# ESCRT-0 regulates AMPA receptor currents and Ca^2+^- dependent signaling

**DOI:** 10.64898/2026.06.19.733273

**Authors:** Mahsa Pourhamzeh, Lara Dozier, Amanda Wilpitz, Yixing Du, Daniel B. McClatchy, Mariel K. B. Micael, Joshua E. Mayfield, Stephen K. Gilmore-Hall, Josephina E. Ronson, Katrin Soldau, Donald P. Pizzo, Brent Aulston, Erin E. Sullivan, Timothy F. Shay, Jin Wang, Subhojit Roy, Viviana Gradinaru, Justin H. Trotter, Kim Dore, John R. Yates, Gentry N. Patrick, Christina J. Sigurdson

**Author notes:** Department of Biochemistry & Molecular Biology, SUNY Upstate Medical University, Syracuse, NY, 13210. Corresponding authors: Christina J. Sigurdson, Department of Pathology, UC San Diego, 9500 Gilman Dr., La Jolla, CA 92093, USA; Gentry N. Patrick, Department of Neurobiology, UC San Diego, 9500 Gilman Dr., La Jolla, CA 92093, USA. **Abbreviated title**: ESCRT-0 function at the post-synapse. **Competing interests:** The authors declare no competing financial interests.

## Abstract

Membrane protein trafficking is essential for synaptic growth, maintenance, function, and plasticity, and involves the regulated exocytosis and endocytosis of proteins to and from the pre-and post-synaptic membranes. Defects in the clearance of membrane proteins can lead to the accumulation of ubiquitinated membrane proteins and contribute to neurodegenerative disease. The ESCRT (endosomal sorting complexes required for transport) machinery binds and sorts ubiquitinated membrane proteins into lysosomes for degradation, yet the presence and function of ESCRTs in sorting ubiquitinated AMPA and other receptors at the post-synapse remain unclear. Here we show that the ubiquitin-binding ESCRT-0 protein, Hrs, localizes to both pre- and post-synapses, and levels are modulated by neuronal activity, increasing and decreasing with higher and lower neuronal activity, respectively. Phosphoproteomic profiling of Hrs-depleted post-synaptic membranes revealed a role for Hrs in glutamatergic synaptic transmission, including long-term potentiation. In addition, Hrs-depleted neurons showed faster AMPAR current kinetics and reduced amplitude in whole-cell patch-clamp recordings. Genetic deletion of neuronal *Hgs* in mice led to reductions in phosphorylated CaMKII-α and -β (T286/T287) and structural proteins, PSD-95 and gephyrin, suggestive of LTD (long-term depression)-like synaptic depression. In contrast, Hrs overexpression led to increases in Ca^2+^-dependent signaling, including protein kinase C (PKC) and PKC substrate, AMPAR subunit GluA1-S831, a site which increases conductance. Together, these findings identify a dynamic, bidirectional role for Hrs at the post-synapse as it both senses and is modulated by neuronal activity, ultimately impacting excitatory synaptic strength.

**Significance Statement:** Synaptic plasticity relies on dynamic trafficking and turnover of membrane proteins, including AMPA-type glutamate receptors (AMPARs), yet how receptor trafficking intersects with ubiquitin-mediated sorting pathways at synapses remains unclear. We show that the ubiquitin-binding ESCRT-0 protein, Hrs, localizes to both pre- and post-synapses, and its abundance is bidirectionally regulated by neuronal activity. Genetic depletion of Hrs in mice reduces CaMKII phosphorylation and impacts AMPAR channel surface localization. In contrast, neuronal-specific Hrs overexpression led to enhanced GluA1 and protein kinase C substrate phosphorylation, suggesting altered AMPAR trafficking, subunit composition, and/or function. Thus, Hrs emerges as a modulator of glutamatergic signaling, coupling ubiquitin-mediated receptor sorting to the fine-tuning of synaptic transmission, with direct implications for learning and memory in health and disease.

## Introduction

In neurons, the rapid turnover of synaptic vesicles and post-synaptic receptors is critical to regulate signal transduction and maintain homeostatic synaptic plasticity (Ehlers, 2000; Rizzoli, 2014). Synaptic transmembrane proteins tagged with ubiquitin are endocytosed and transit into a tightly regulated process for degradation by proteasomes or ESCRT-mediated delivery to lysosomes (Schwarz and Patrick, 2012; Alvarez-Castelao and Schuman, 2015). At both pre- and post-synaptic terminals, proteasomes selectively degrade proteins tagged with polyubiquitin chains, whereas ESCRT machinery targets mono- or short-chain ubiquitinated transmembrane proteins for lysosomal degradation (Jarome and Helmstetter, 2013; Soykan et al., 2021). While ESCRT activity has been described in axons, its function and regulation in mediating synaptic receptor trafficking at the post-synapse are incompletely understood. Altered internalization, endocytic sorting, or degradation of synaptic proteins would be expected to cause an accumulation of ubiquitinated proteins and disrupt proteostasis, a hallmark of neurodegenerative disease (Hipp et al., 2019; Hetz, 2021). Identifying the physiologic trafficking mechanisms for post-synaptic receptor clearance is therefore a critical component in understanding synapse dysfunction during disease.

The ESCRT system consists of four complexes (ESCRT-0, -I, -II, and -III) that act sequentially to sort endocytosed ubiquitinated membrane proteins into intraluminal vesicles (ILVs) within multivesicular bodies (MVBs) for lysosomal degradation or exosomal release (Hurley et al., 2025). The binding of monoubiquitinated proteins is initiated by ESCRT-0, composed of hepatocyte growth factor-regulated tyrosine kinase substrate (Hrs) and signal transducing adapter molecule 1 (STAM1), with Hrs serving as the primary gatekeeper for ubiquitinated cargo (Bache et al., 2003; Ren and Hurley, 2010). Hrs exhibits more than twice the affinity for ubiquitin as compared to STAM1 (Mayers et al., 2011). Mutations within the double ubiquitin-interacting motif (DUIM) domain of Hrs abolish the ability of ESCRT-0 to form stable complexes with ubiquitinated cargo (Hirano et al., 2006; Mayers et al., 2011).

*Hgs* mRNA is highly expressed in brain regions crucial for learning and memory, including the hippocampus, cerebral cortex, and hypothalamus (Tamai et al., 2008). At pre-synaptic terminals, Hrs contributes to synaptic vesicle (SV) degradation and undergoes activity-dependent re-distribution (Birdsall et al., 2022). Increased neuronal firing enhances Hrs accumulation in axons and SV pools, suggesting a direct link to neuronal activity (Sheehan et al., 2016; Birdsall et al., 2022). Furthermore, a spontaneous point mutation in the Hrs gene (*Hgs*) in “teetering” mice (Hrs*^tn/tn^*) causes neuromuscular weakness, synaptic dysfunction, and disrupted neurotransmitter release at the neuromuscular junction (Watson et al., 2015), suggestive of an essential role for ESCRT-0 in synaptic processes. Hrs also modulates epidermal growth factor receptor (EGFR) recycling and autophagic flux, indicative of its broader role in supporting neuronal homeostasis (Stern et al., 2007).

We previously reported a marked reduction of Hrs and STAM1 in both human and experimental prion disease (Lawrence et al., 2023). Neuronal deletion of *Hgs* accelerated the accumulation of ubiquitinated proteins at synapses, exacerbated synaptic pathology, and reduced survival time in mice. Moreover, prion-infected mice lacking neuronal Hrs also showed a faster accumulation of hyperphosphorylated AMPA receptor (AMPAR) subunit GluA1 at S845 and S831, modifications that increase surface retention and channel conductance, respectively (Derkach et al., 1999; Lee et al., 2000; Diering et al., 2014), raising the question of whether Hrs modulates the trafficking and turnover of AMPAR in health and disease.

Surface AMPARs can become dephosphorylated at specific sites to promote their ubiquitination, internalization, and endocytic sorting to lysosomes, which degrade AMPARs in an activity-dependent manner for plasticity paradigms such as homeostatic downscaling (Schwarz et al., 2010; Scudder et al., 2014; Widagdo et al., 2017). Specifically, the E3 ligase Nedd4-1 ubiquitinates GluA1 at Lys-868 to facilitate AMPAR sorting and degradation, thereby reducing receptor surface expression (Lin et al., 2011). Excess GluA1 ubiquitination has been implicated in amyloid-β (Aβ)-induced removal of synaptic AMPARs associated with dendritic spine loss and synaptic depression in AD (Hsieh et al., 2006; Rodrigues et al., 2016).

Despite the well-established role of ESCRT-0 in the endo-lysosomal pathway, its distribution, regulation by neuronal activity, and specific role at synapses, including the trafficking and turnover of AMPARs, remain largely unknown. Here we show that the ubiquitin-binding ESCRT-0 protein, Hrs, localizes to the pre- and post-synapse, and its abundance is dynamically and bidirectionally regulated by neuronal activity, with increases or decreases in neuronal activity leading to increases or decreases in Hrs, respectively. Moreover, depleting neuronal Hrs in mice impacted AMPAR function and the levels of phosphorylated CaMKII, whereas Hrs overexpression increased protein kinase C phosphorylated substrates and phosphorylated AMPAR subunit (pGluA1-S831) levels, suggesting that Hrs acts as a sensor at the post-synapse to fine-tune neuronal activity.

## Results

### Hrs localizes to pre- and post-synaptic neuronal compartments

The rapid clearance of ubiquitinated membrane receptors at the post-synaptic density (PSD) is critical to synaptic plasticity (Tai and Schuman, 2008). Given that ESCRT-0 binds and sorts ubiquitinated membrane proteins, we first determined the subcellular localization of neuronal Hrs at the pre- and post-synapse. We purified the pre-synapse and PSD from the brain of wild-type (WT) mice (Figure S1) and immunoblotted fractions for synaptophysin and PSD-95 as indicators of the pre-synaptic (supernatant 3; S3) and post-synaptic density (PSD2) fractions, respectively, finding little overlap between S3 and PSD2 (Figure 1A). Hrs was enriched in S3 relative to the initial supernatant fraction, S1 [S3 / S1: 1.42 ± 0.8 (mean ± SEM) versus synaptophysin: 0.87 ± 0.2, and PSD95: 0.42 ± 0.08]. Notably, Hrs was also present in PSD2 (PSD2 / S1: 0.33 ± 0.07 versus synaptophysin: 0.008 ± 0.0, and PSD95: 3.03 ± 1) (Figure 1A-B). CHMP2B (ESCRT-III) was also present in PSD2 (Figure 1A-B), indicating that ESCRT proteins localize to the post-synapse.

**Figure 1.**
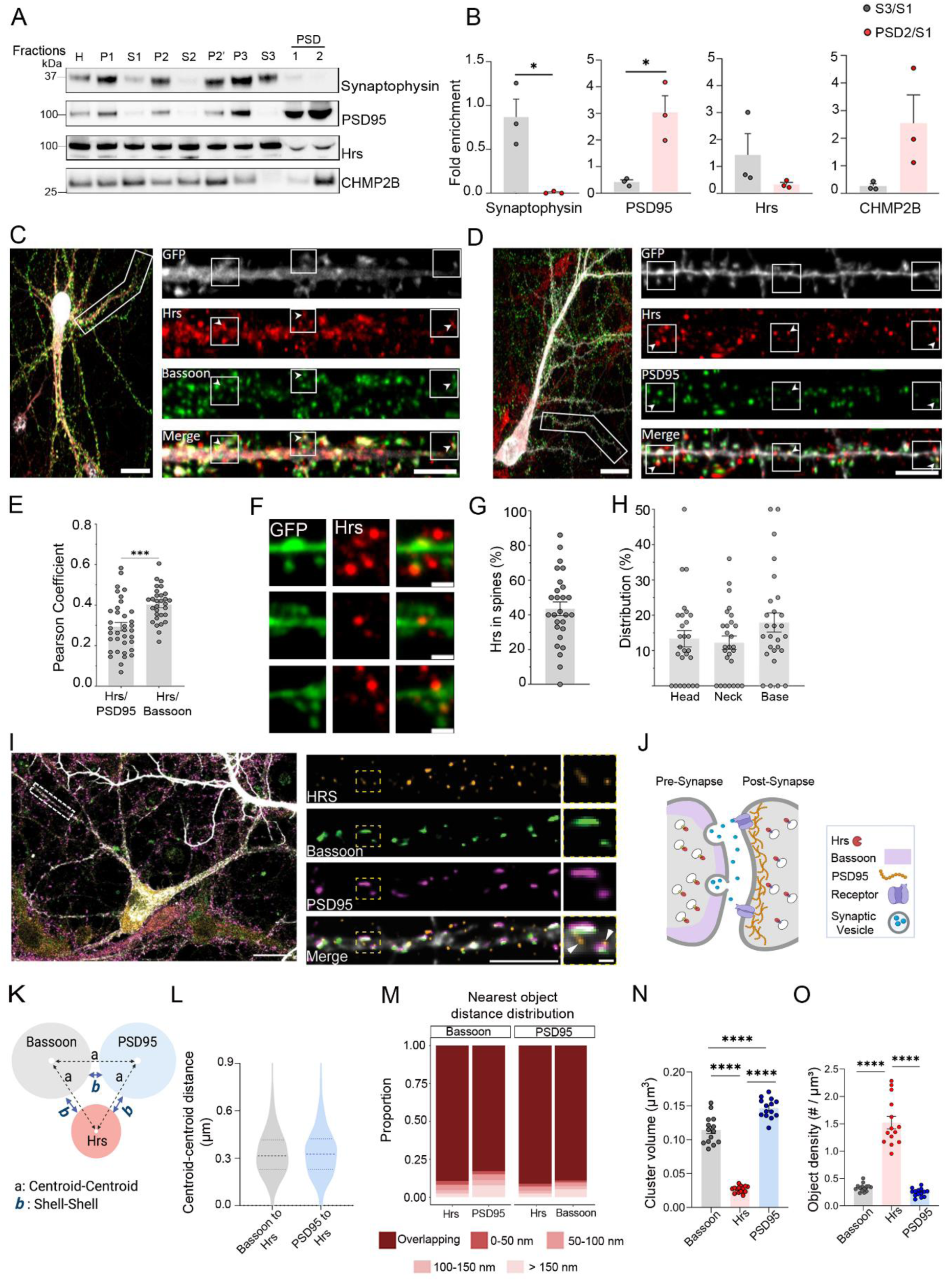
Hrs localizes to the pre- and post-synapse of neurons. (A) Representative western blot and (B) quantification of subcellular fractions probed for synaptophysin, PSD-95, Hrs (ESCRT-0), and CHMP2B (ESCRT-III). Equivalent amounts of protein were loaded into each lane within each experiment. *N* = 3 independent experiments. (C-D) Representative images of rat hippocampal neurons immunolabeled for endogenous Hrs and Bassoon (C) or Hrs and PSD-95 (D) in GFP-filled dendrites (white). (E) Quantification of the Pearson’s correlation coefficients reveals the extent of colocalization between Hrs and Bassoon or Hrs and PSD-95; *n* = 30 and 36 dendrites for Hrs/Bassoon and Hrs/PSD-95, respectively, from 2 (Hrs/Bassoon) and 3 (Hrs/PSD-95) independent experiments. (F) Representative high magnification images of neurons (boxed region in panel (D) immunolabeled for endogenous Hrs (red) and PSD-95 (green) show Hrs within single spines. (G) Quantification of spines harboring Hrs (*n* = 320 spines). (H) Of the spines with Hrs, the distribution to head, neck, or base of spines was quantified; 27 dendritic segments were analyzed from 2 independent experiments. (I) Representative super-resolution image of cultured neurons immunolabeled for Hrs (orange), Bassoon (green), and PSD95 (magenta). Higher-magnification views of the boxed region indicate Hrs-positive structures within dendritic spines (arrowheads). N = 3 independent cultures with 10 neurons imaged per culture. (J) Schematic representation illustrates the localization of Hrs relative to Bassoon and PSD95, with Hrs positioned in close proximity to pre- and postsynaptic compartments. (K) Diagram depicts the distance measurements used for nanoscale spatial analysis. Distances were measured from centroid-to-centroid (a) or from shell-to-shell. (L) Violin plots show the centroid-to-centroid distances between Hrs and Bassoon and between Hrs and PSD95. Dashed lines indicate median and quartile distributions. (M) Distribution graph shows the nearest object distances between Hrs, Bassoon, and PSD95 clusters (based on shell-to-shell distances). Stacked bars represent the proportion of objects (proteins) that overlap or are within defined distance bins (0–50 nm, 50–100 nm, 100–150 nm, and >150 nm), and demonstrates extensive spatial association of Hrs with both synaptic markers. (N) Quantification of cluster volume for Bassoon, Hrs, and PSD95. Each dot represents one image, plotted as the mean cluster volume. (O) Quantification of object density for Bassoon, Hrs, and PSD95 (# / µm^3^). Each dot represents the average object density per image. Scale bars (C-D) = 20 and 5 µm for whole cell and dendrite images, respectively. Scale bars (I) left panel = 5 µm (dendrite) and right panel = 200 nm (synapse). Mean ± SEM (B, E, G, H, L, N, and O). Unpaired two-tailed *t*-test (B, E, and L) and one-way ANOVA with Tukey’s multiple comparison test (N and O); *\* p <* 0.05; *** *p <* 0.001, and **** *p* < 0.0001.

To further validate and quantify Hrs synaptic localization, we immunolabeled hippocampal neurons for Hrs, Bassoon, and PSD-95, and quantified their spatial overlap. Hrs co-localized with pre-and post-synaptic proteins, and was more abundant at the pre-than the post-synapse (Pearson coefficient (mean ± SEM): 0.40 ± 0.02 and 0.29 ± 0.02 for Hrs/Bassoon and Hrs/PSD-95, respectively) (Figure 1C-E). Hrs localized to approximately 40% of dendritic spines and was within the head, neck, and base (Figure 1F-H), suggesting that within Hrs-positive spines, Hrs is distributed throughout the spine.

To define the nanoscale distribution of Hrs at synapses, we immunolabeled hippocampal neurons for Hrs, Bassoon, and PSD-95 and performed super-resolution microscopy at the synapse using stimulated emission depletion (3D-STED). Bassoon and PSD-95 were closely associated with Hrs nanodomains, frequently overlapping (89-91%) (Figure 1I-M and Figure S2A), however Hrs clusters were smaller and denser than Bassoon or PSD95 clusters (Figure 1N-O). Notably, the distance between the edges of Hrs and Bassoon versus Hrs and PSD-95 were similar (Figure S2B–G). Thus, Hrs is positioned within nanoclusters to function near the pre-synaptic active zone and post-synaptic compartments.

### Synaptic activity regulates Hrs levels

Given that Hrs localizes to the post-synapse, we next tested whether alterations in neuronal activity modulate Hrs levels at the post-synapse. Prolonged changes in neuronal activity induce synaptic plasticity pathways, including homeostatic mechanisms that upscale or downscale activity, in part by the insertion or removal of AMPARs (Pozo and Goda, 2010; Davis, 2013). To determine whether Hrs expression can be modulated by synaptic activity, we exposed rat cortical neurons to either bicuculline (Bic; 20 μM; GABA-A receptor antagonist) or tetrodotoxin (TTX; 2 μM; sodium channel blocker) to increase or reduce neuronal activity, respectively, or vehicle, for 72 hours. Surprisingly, Hrs increased by 34% in Bic-treated neurons [1.54 ± 0.17 versus 1.15 ± 0.03 (untreated); *p* < 0.05] and decreased by 31% in TTX-treated neurons [0.80 ± 0.12 versus 1.15 ± 0.03 (untreated); *p* < 0.05] (Figure 2A-B), indicating that Hrs protein levels are dynamically regulated by neuronal activity in a manner similar to phosphorylated AMPARs. We further confirmed the effect of Bic on Hrs using immunofluorescence labeling of endogenous Hrs in hippocampal neurons. Bic treatment resulted in a 22% increase in Hrs labeling when compared to untreated controls (1.22 ± 0.06 versus 0.98 ± 0.05 for Bic and control, respectively; *p* < 0.05) (Figure 2C-D).

**Figure 2.**
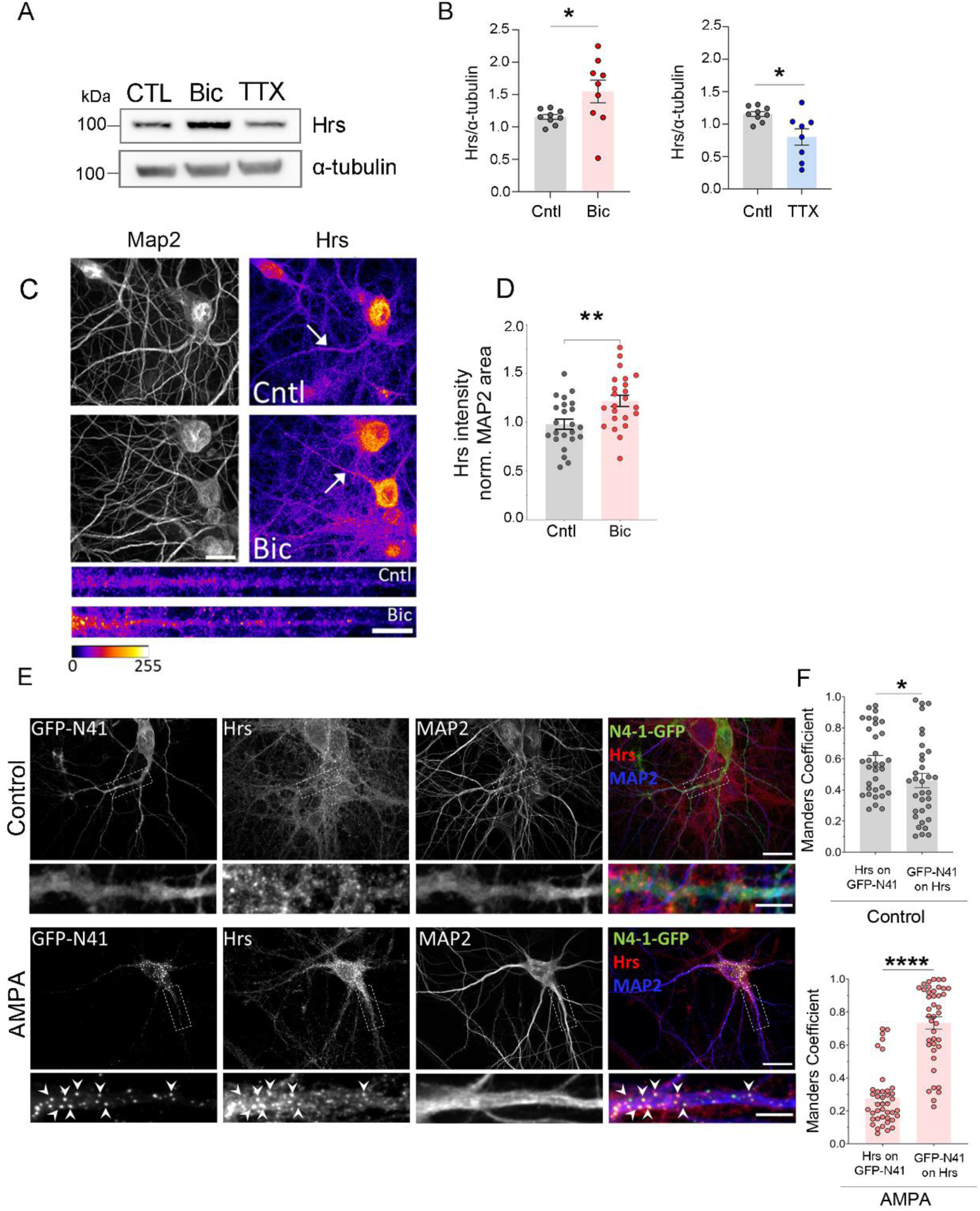
Hrs expression is dynamically regulated by neuronal activity. Rat cortical neurons were treated with vehicle, bicuculline (Bic, 20 μM), or tetrodotoxin (TTX, 2 μM) for 72 hours. (A) Representative western blot and (B) quantification of Hrs and α-tubulin (loading control). *N* = 8 independent experiments. (C) Representative images of hippocampal neurons treated with Bic (20 μM; 72 hours) or vehicle and immunolabeled for endogenous Hrs (red) and MAP2 (blue). (D) Quantification of Hrs intensity normalized to MAP2 area (binary mask); *n* = 23 fields of neurons per condition from 2 independent experiments. (E) Representative images of hippocampal neurons expressing Nedd4-1-GFP (N4-1-GFP; green) treated with AMPA (10 μM; 10 minutes) or vehicle control and immunolabeled for Hrs (red) and MAP2 (blue). (F) Quantification of Manders colocalization coefficient with thresholding to assess overlap between Hrs with GFP-Nedd4-1 with and without AMPA treatment. *N* = 34 and 41 dendrites analyzed for control and AMPA-treated neurons from 3 independent experiments. Arrowheads indicate N4-1-GFP and Hrs puncta, which co-localize following AMPA treatment. Scale bars = 20 µm and 5 µm for whole-cell and dendrite images, respectively. Data are presented as mean ± SEM. Unpaired two-tailed Student’s *t*-test test (panels B, D, and F). * *p* < 0.05, ** *p* < 0.01, **** *p* < 0.0001.

### E3 ubiquitin ligase, Nedd4-1, co-localizes with Hrs in response to AMPA

GluA1-containing AMPARs are rapidly ubiquitinated by the E3 ubiquitin ligase, Nedd4-1, in response to direct activation (Schwarz et al., 2010). To next determine whether AMPA receptor activation promotes an association between Hrs and the E3 ubiquitin ligase, Nedd4-1, we treated primary hippocampal neurons expressing GFP-tagged Nedd4-1 with AMPA (20 μM) or vehicle control for 10 minutes, and immunolabeled cells for endogenous Hrs and MAP2. GFP-Nedd4-1 redistributed from a diffuse to a punctate pattern in both somatic and dendritic compartments following AMPA treatment, similar to what was previously shown for HA-tagged Nedd4-1 constructs (Scudder et al., 2014). Notably, we found a marked overlap between Hrs and GFP-Nedd4-1 in the AMPA-treated group as compared to control-treated neurons (Figure 2E-F) (*p* < 0.05), suggestive of AMPA-induced Nedd4-1 recruitment to ESCRT-0 endosomal compartments.

### Hrs levels impact AMPAR kinetics in CA1 pyramidal neurons

Since Hrs protein levels are dynamically regulated by neuronal activity, we next assessed the functional effects of reducing Hrs on excitatory synaptic transmission. We performed whole-cell patch clamp recordings from CA1 pyramidal neurons in organotypic hippocampal slices using AAV to mediate depletion of Hrs via shRNA (Figure 3A). Effective knockdown of Hrs was confirmed by reduced immunofluorescence intensity in hippocampal neurons (Figure S3). Notably, Hrs knockdown (KD) neurons had a lower percentage of cells with slow AMPAR EPSCs (21 out of 41 recorded neurons) compared to scrambled control neurons (38 out of 46 recorded neurons; *p* = 0.002) (Figure 3B-F). Each trace represents averaged sweeps from a single neuron; the slow, multi-peak profile was a reproducible feature of those neurons rather than a stochastic artifact (Figure 3C). Representative AMPAR and NMDAR currents recorded from the same neurons, expressing either the scrambled control or the Hrs shRNA, are shown in Figure S4. As these slow AMPAR currents are biologically distinct responses and differ between groups, they were quantified rather than excluded (Figure 3C). Importantly, the AMPA/NMDA ratio was reduced in *Hrs* KD neurons with current measured at 100 ms post-stimulation (Figure 3G-H), which may reflect a relative reduction in the strength or number of AMPAR-mediated synaptic responses (Béïque et al., 2006; Counotte et al., 2014). This could arise from a decrease in either the number or function of AMPARs, leading to weaker fast excitatory synaptic transmission, which would result in a reduced initial depolarizing current. Alternatively, an increased NMDAR-mediated response or function can also lower the AMPA/NMDA ratio (Fourgeaud et al., 2010). To further examine the impact on AMPAR current dynamics, we analyzed the rise time and decay slope of AMPAR EPSCs (Figure 3I-L). EPSCs were aligned to the point of maximal rise prior to kinetic analysis, and recordings were monitored to ensure stable access resistance across groups. Hrs-depleted neurons displayed a decrease in AMPAR rise time (Figure 3I), indicating faster receptor activation kinetics and more rapid synaptic signal initiation and transmission (Kleppe and Robinson, 1999; Koike-Tani et al., 2005; Jacobi and von Engelhardt, 2021). Additionally, we found an increased AMPAR decay slope in Hrs-depleted neurons (Figure 3L), suggesting a faster decline or shorter duration of the excitatory post-synaptic current following receptor activation (Joshi et al., 2004), consistent with the lower percentage of slow AMPAR currents. Loss of Hrs-dependent endosomal sorting may alter AMPAR subunit composition and/or association with auxiliary subunits (such as transmembrane AMPA receptor regulatory proteins, TARPs), resulting in AMPARs with faster deactivation and desensitization kinetics. Together, these findings suggest that Hrs contributes to the regulation of AMPAR kinetics and maintenance of excitatory synaptic inputs in CA1 pyramidal neurons.

**Figure 3.**
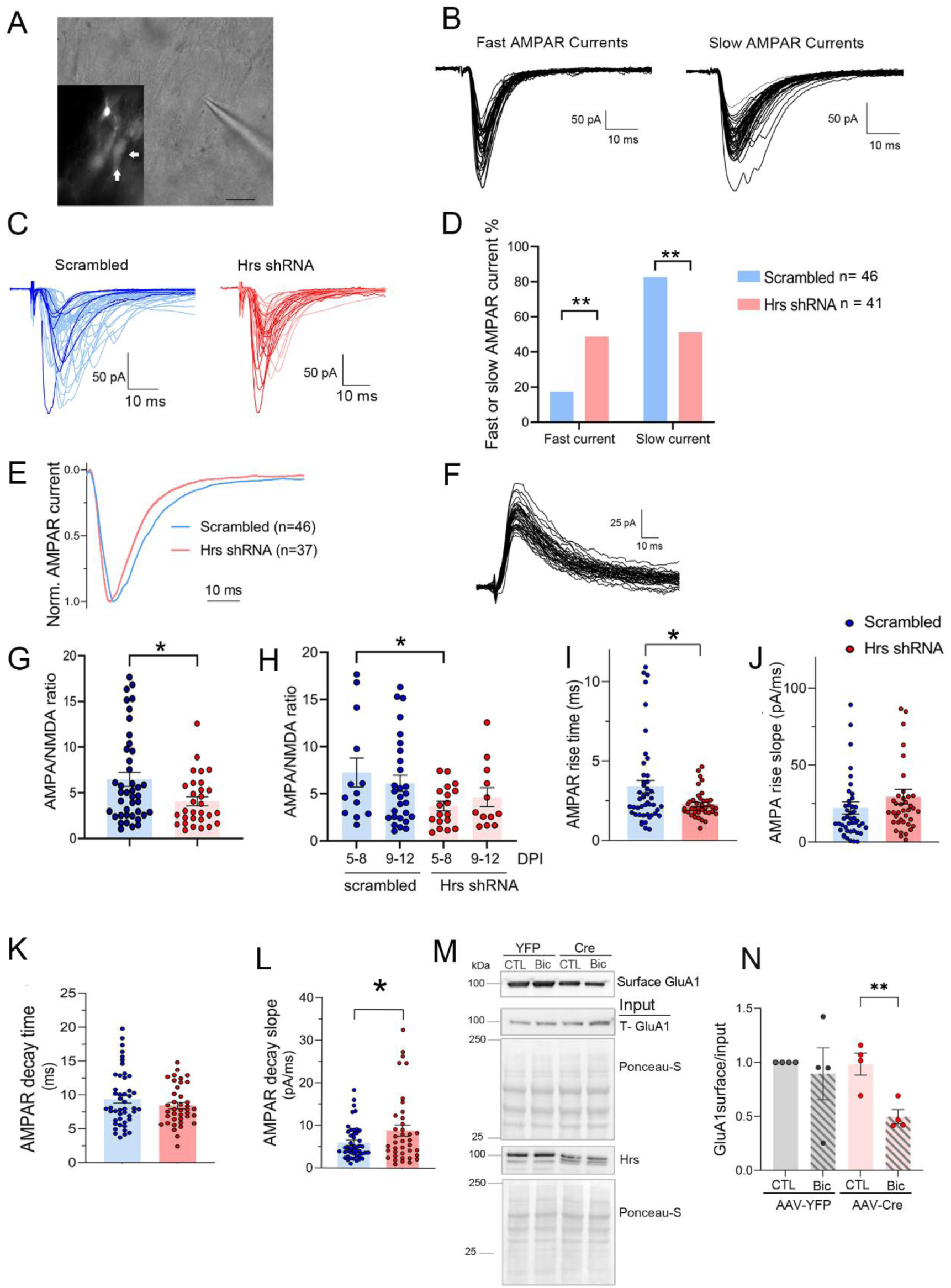
Depletion of Hrs reduces the AMPA/NMDA ratio and alters AMPAR kinetics in CA1 pyramidal neurons. (A) Patch-clamp recorded hippocampal CA1 pyramidal neuron in an organotypic slice. The GFP fluorescence (inset) indicates Hrs knockdown (arrows). (B) Two types of AMPAR EPSCs, fast and slow, were observed in both scrambled and Hrs knockdown neurons. (C) Recording traces of AMPAR EPSCs from scrambled and Hrs knockdown neurons. Each trace represents the average current recorded from each neuron. Dark blue/red: fast AMPAR EPSCs; light blue/red: slow AMPAR EPSCs. (D) Hrs-depleted neurons show a lower percentage of cells with slow AMPAR EPSCs and a higher percentage of cells with fast AMPAR EPSCs. (E) Averaged normalized AMPAR EPSCs of scrambled control and Hrs knockdown neurons. Traces were aligned by the point of maximal rise to minimize variability in evoked current latency during kinetic analysis. (F) Representative recording traces of NMDAR currents recorded at +40 mV. (G) Hrs knockdown reduced the AMPA/NMDA ratio, with NMDA currents measured at 100 ms after stimulation. (H) AMPA/NMDA ratio analyzed in slices at 5–8 or 9–12 days post-infection (DPI), using NMDAR currents measured at 100 ms after stimulation. Recordings grouped by 5–8 or 9–12 DPI showed the same AMPA/NMDA ratio, indicating that efficient knockdown was achieved by 5 DPI and DPI did not affect the AMPA/NMDA ratio in either scrambled or Hrs knockdown neurons. Hrs knockdown reduced the AMPA/NMDA ratio in both groups, indicating that efficient knockdown is achieved by 5 DPI. (I–L) Quantification of AMPAR EPSC rise time, decay time, and slope from scrambled control and Hrs knockdown neurons. (M) Representative immunoblots of surface biotinylated GluA1 and input fractions from *Hrs^f/f^* cortical neurons expressing control AAV-YFP or AAV-Cre treated with DMSO or bicuculline (Bic, 20 μM) for 24 h. Input samples were probed for total GluA1, Hrs, and Ponceau S as a loading control. (N) Quantification of surface GluA1 normalized to input GluA1 (each dot represents one biological replicate). Data are presented as mean ± SEM and assessed by Chi-square (Fisher’s exact) test for panel D, or unpaired, two-tailed Student’s t-tests for panels G–L, and N; * *p* < 0.05, ** *p* < 0.01.

### Hrs impacts activity-dependent surface trafficking of GluA1

Given the alterations in AMPAR-mediated currents observed following Hrs depletion, we next assessed whether Hrs can alter the surface abundance of AMPA receptors. We ablated Hrs from Hrs^f/f^ cortical neurons and measured surface GluA1 under basal conditions and following Bic-treatment for 24 hours. Under basal conditions, Cre-mediated Hrs depletion did not alter surface GluA1 relative to controls (Figure 3M-N), indicating that Hrs is not required for maintenance of steady-state surface GluA1 levels. However, following Bic-induced neuronal activation, Hrs-ablated neurons exhibited an approximately 40% reduction in surface GluA1 compared with Bic-treated control neurons (Cre + Bic: 50 ± 7%; YFP-Bic: 90 ± 24%; *p* < 0.01), suggesting that Hrs impacts the activity-dependent regulation of surface GluA1.

### Depleting Hrs disrupts glutamatergic signaling at the post-synaptic membrane

Activity-dependent phosphorylation and dephosphorylation events play a significant and in many cases rate-limiting role at synapses to control the insertion, retention, and internalization of membrane proteins, including AMPA receptors, at the post-synaptic membrane (Swope et al., 1999). To test whether Hrs impacts the recycling of synaptic proteins by modulating local kinase and phosphatase activity, we isolated the post-synaptic membranes from the cortices of neuronal Hrs-depleted (Tamai et al., 2008) and control mice, and performed phosphoproteomics profiling (*Hrs^f/f^Syn1-Cre^+/-^*, *n* = 4; *Hrs^f/f^Syn1-Cre^-/-^*, n = 5). Peptides were tandem mass tag (TMT) labeled followed by mass spectrometry to quantitatively profile phosphopeptides (Figure 4A). Proteins with a *p*-value less than 0.05 were considered differentially expressed. Phosphopeptide profiles clearly separated *Hrs^f/f^Syn1-Cre^-/-^* and *-Cre^+/-^* groups, with strong reproducibility among replicates (Figure 4B). A total of 924 unique phosphopeptides were identified, with 16 showing differential abundance (please see Supplementary Data 1 for full dataset). Of these, 14 were hyper-phosphorylated, while two were hypo-phosphorylated.

**Figure 4.**
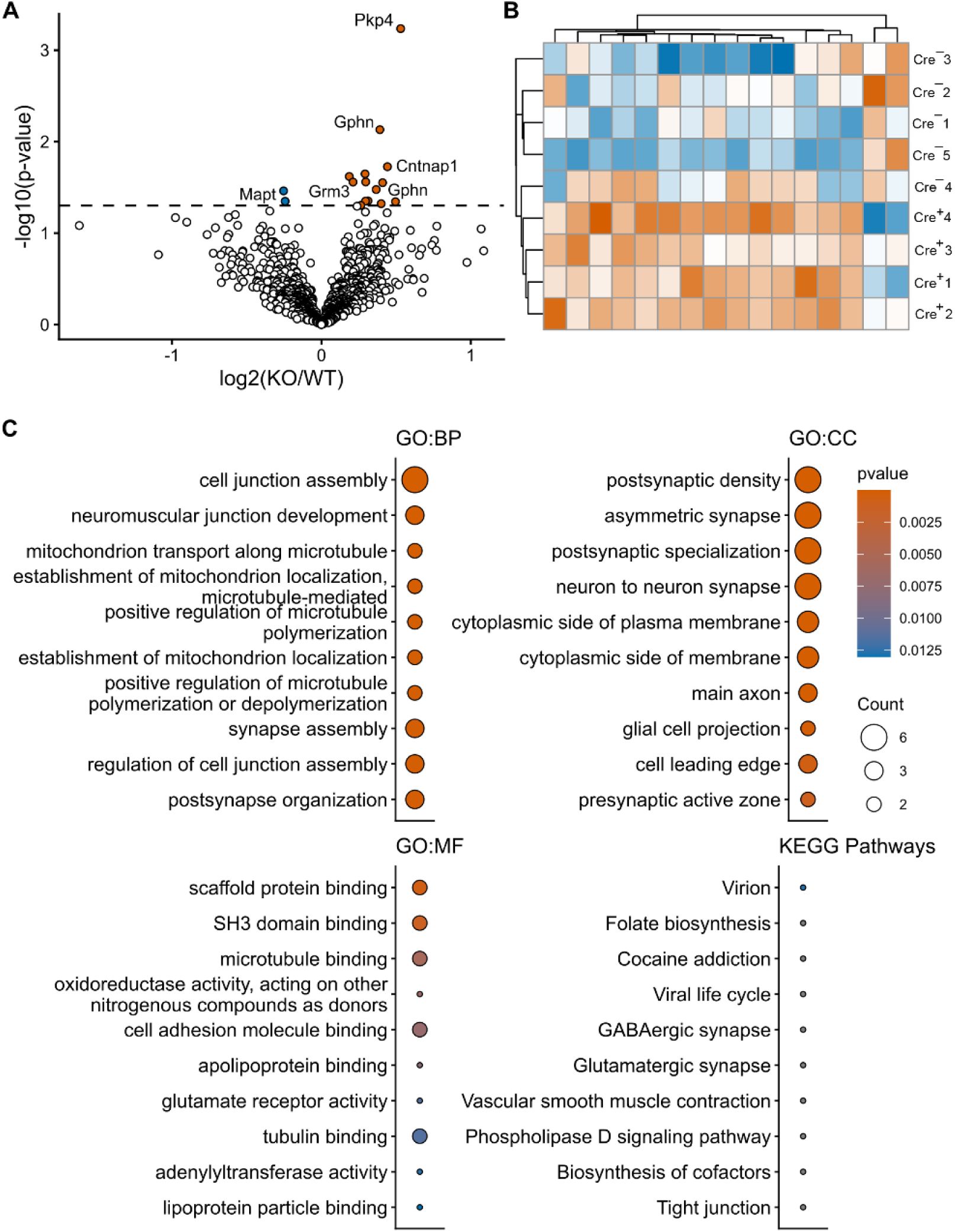
Phosphopeptide abundance and functional enrichment in Hrs-depleted post-synaptic membrane from mouse cortex and hippocampus. (A) Volcano plot shows the differentially abundant phosphopeptides from *Hrs^f/f^Syn1-Cre^+/-^* and *-Cre^-/-^* post-synaptic membrane. The dashed horizontal line indicates the significance threshold at –log10 (0.05). (B) Heatmap of differentially abundant phosphopeptides with hierarchical clustering of samples. Color gradient indicates scaled relative phosphopeptide abundance (vermilion = high, blue = low). Sample numbers (Cre^-^ 1 - 5 and Cre^+^ 1 - 4) correspond to the clustering order. (C) Gene ontology (GO) enrichment analysis of differentially abundant phosphopeptides, showing top biological process (BP), cellular component (CC), and molecular function (MF) categories. KEGG is used in enrichment analyses to identify pathways related to differentially abundant phosphopeptides. The color gradient indicates unadjusted *p*-values. Point size corresponds to the number of genes enriched within a given ontology/pathway.

Gene Ontology (GO) analyses revealed that phosphoproteins are involved in biological processes, including cell junction assembly, synapse organization, and post-synaptic structure (Figure 4C). The cellular compartment ontology further supported the localization of these differentially phosphorylated proteins to the PSD and pre-synaptic active zones, as well as glial cell projections (Figure 4C). Functionally, differentially phosphorylated proteins exhibit diverse roles, ranging from scaffold protein interactions and microtubule or tubulin binding to glutamate receptor activity. Kyoto Encyclopedia of Genes and Genomes (KEGG) pathway analysis unified these ontologies by highlighting statistically significant alterations in GABAergic and glutamatergic synapses, as well as tight junctions (Figure 4C). Notable phosphopeptide modifications were identified in key synaptic and cytoskeletal proteins, including hyperphosphorylated contactin-associated protein 1 (Cntnap1; S1372 and S1385), gephyrin (S264 and S268), glutamate metabotropic receptor 3 (Grm3; S878), R3H domain-containing protein 2 (R3hdm2; S347), and plakophilin-4 (Pkp4; S336). Tau (Mapt) was hypophosphorylated at S688, S692, and S696, which are sites linked to neuronal activity and are hyperphosphorylated in AD (Mast et al., 2021). Microtubule-associated protein 1B was also hypophosphorylated (Map1b; S1151, S1153). Collectively, the integrated analysis demonstrates that Hrs-depletion alters proteins involved in cellular and subcellular organization as well as in response to key neuronal signaling molecules such as GABA and glutamate, highlighting the broad impact of Hrs on synaptic structure and signaling dynamics. To complement the phosphoproteomic analysis, total proteomic profiling was also conducted and revealed alterations in ribosomal subunits and translation at the pre- and post-synapse (Figure S5).

### Depleting Hrs from neurons reduces phosphorylated CaMKII-α and -β at T286 and T287, but increases total CaMKII in the cortex

Since Hrs expression can be controlled by neuronal activity, we next examined whether manipulating Hrs expression in neurons affects Ca^2+^ signaling, including key plasticity kinase pathways and scaffolding proteins. We assessed the cortex from adult *Hrs^f/f^Syn1-Cre^+/-^* and *Cre^-/-^* mice and found Hrs reduced in *Cre^+/-^* mice by 53% compared to *Cre^-/-^* controls (0.47 ± 0.07, mean ± SEM) (Figure 5).

**Figure 5.**
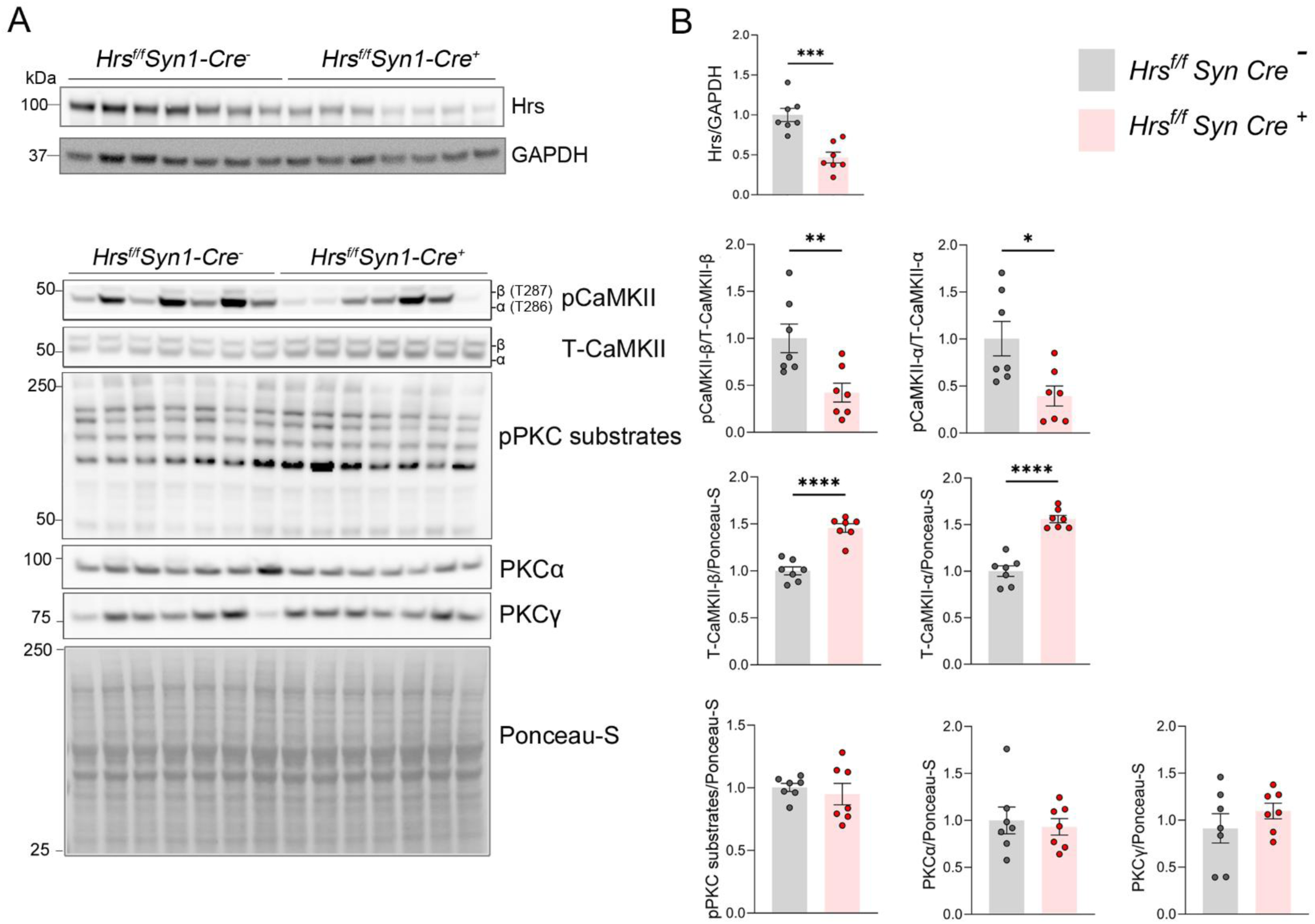
Mice with neuronal Hrs depletion show increased total CaMKII. (A) Western blots of cortical lysates from *Hrs^f/f^Syn1-Cre^-/-^* and *Hrs^f/f^Syn1-Cre^+/-^*mice (*n* = 7 per group). (B) Densitometric quantification of western blots. Mean ± SEM. Unpaired two-tailed Student’s *t*-test; * *p* < 0.05, ** *p* < 0.01, **** *p* < 0.0001.

To determine whether Hrs depletion affects activity-dependent kinase signaling, we measured calcium/calmodulin-dependent protein kinase II (CaMKII) activity through phosphorylation (p) at T286 and T287 in cerebral cortical lysates. pCaMKII-α and -β were markedly reduced (60% decrease: 0.39 ± 0.10 and 0.42 ± 0.10, respectively) (Figure 5), suggestive of reduced activity (Lučić et al., 2023). Strikingly, total CaMKII-α and -β were increased by 56% (1.56 ± 0.03) and 46% (1.45 ± 0.04), respectively (Figure 5). The decrease in pCaMKII and increase in total CaMKII levels suggest a neuronal activity state with markedly altered CaMKII activity. Interestingly, other Ca^2+^-sensitive kinases and their substrates and phosphatases, including phosphorylated-PKA (pPKA) and -PKC (pPKC) substrates, PKCα and γ (Jenkins and Traynelis, 2012; Diering and Huganir, 2018; Summers et al., 2019), and PP2B (calcineurin), were unchanged (Figure 5 and Figure S6), suggesting that Hrs depletion does not broadly affect PKA-, PKC-, or PP2B-mediated signaling pathways. These data suggest that loss of Hrs leads to a specific reduction of CaMKII autophosphorylation, suggestive of reduced neuronal activity.

To assess whether Hrs depletion affects synaptic architecture, we examined key scaffolding proteins involved in inhibitory and excitatory synapses. Western blot analysis revealed a 12% reduction in gephyrin (0.88 ± 0.01) and a 17% reduction in PSD-95 (0.83 ± 0.05) in *Hrs^f/f^Syn1-Cre^+/-^*mice relative to controls (Figure 6), suggestive of a reduction in scaffolding proteins at the PSD. The presynaptic protein, synaptophysin, was unaffected. We did not observe any changes in NMDARs, however for AMPARs, there was a modest increase in pGluA1-S831 (Figures S7 and S8). The reduction in PSD-95, coupled with the reduced pCaMKII kinase, suggests that Hrs depletion may lead to impaired synaptic organization.

**Figure 6.**
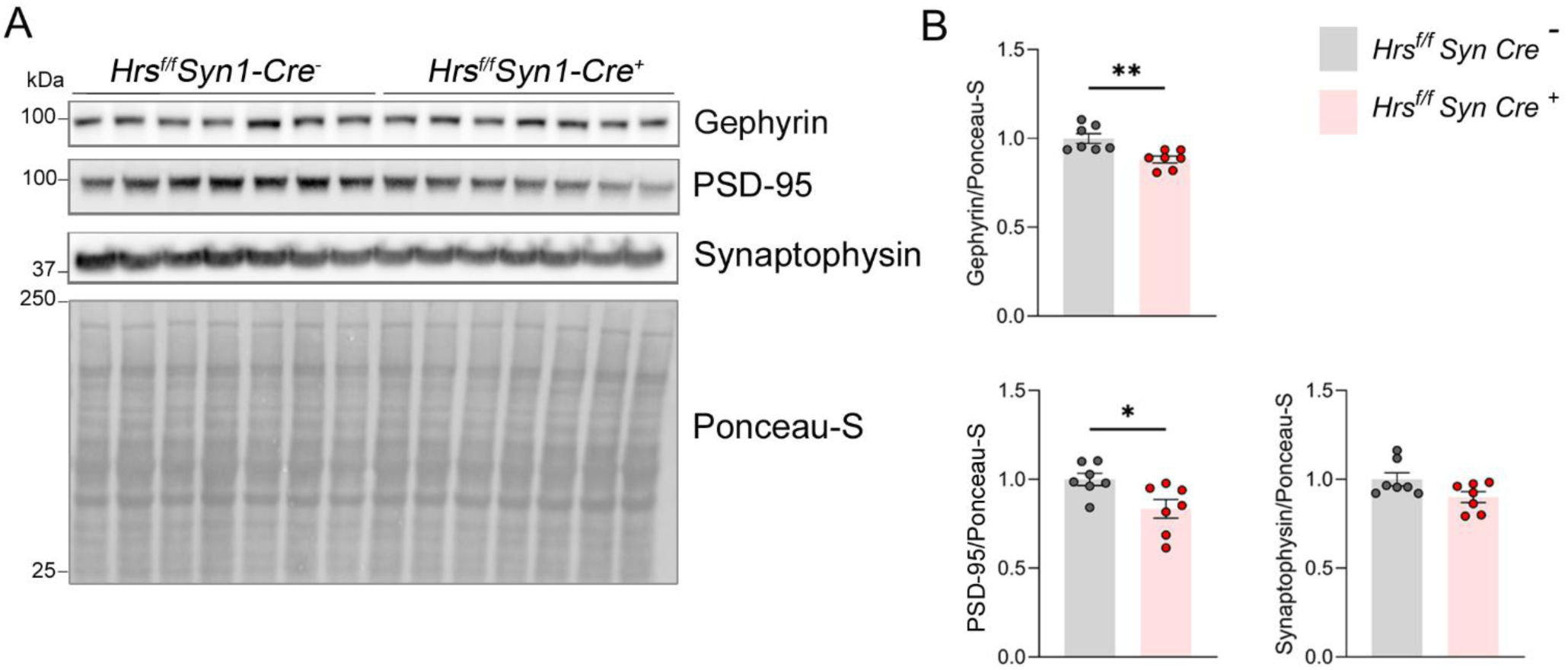
Mice with neuronal Hrs depletion show reduced synaptic scaffolding proteins gephyrin and PSD-95. (A) Western blots of cortical lysates from *Hrs^f/f^Syn1-Cre^-/-^* and *Hrs^f/f^Syn1-Cre^+/-^* mice (n = 7 per group). (B) Densitometric quantification of western blots. Mean ± SEM. Unpaired two-tailed Student’s *t*-test; * *p* < 0.05 and ** *p* < 0.01.

### Neuronal Hrs overexpression enhances PKC activity and phosphorylated GluA1-S831, a PKC substrate site

To investigate the impact of neuronal Hrs overexpression on glutamate receptor regulation, we generated AAV.CAP-B10 vectors (Goertsen et al., 2022) encoding either HA-tagged Hrs or GFP under the hSyn1 promoter and retro-orbitally administered 1.11 × 10¹² viral genomes (vg) of AAV-Hrs-HA or AAV-GFP to adult WT mice. Three weeks post-injection, immunolabeling confirmed robust widespread neuronal Hrs overexpression (Figure S9), which was further validated by western blot on cortical homogenates, showing a 2.7-fold increase in Hrs (Figure 7A-B). Hrs upregulation reportedly promotes ubiquitin-dependent processes that may influence synaptic vesicle protein trafficking and turnover (Birdsall et al., 2022), driving a more rapid or extensive tagging of synaptic proteins for trafficking and subsequent degradation. Notably, ubiquitinated proteins were modestly increased with Hrs overexpression (Figure 7A-B).

**Figure 7.**
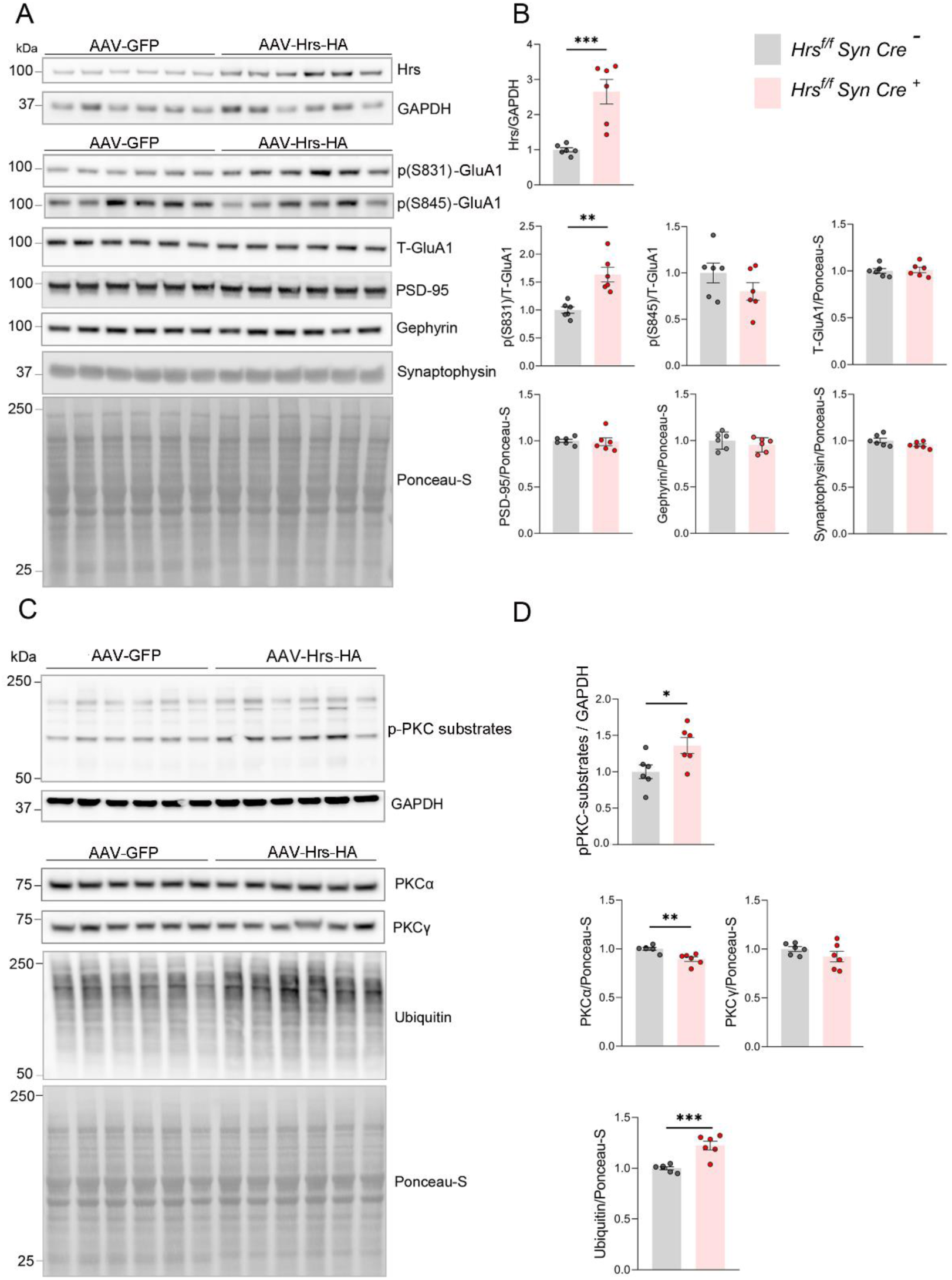
Neuronal Hrs overexpression in mice leads to increases in phosphorylated AMPAR subunit GluA1(S831) and PKC phosphorylated substrates. (A and C) Western blots of cortical lysates from adult WT mice transduced with AAV-GFP or AAV-Hrs-HA (n = 6 per group). (B and D) Densitometric quantification of western blots. Mean ± SEM. Unpaired two-tailed Student’s *t*-test; *p < 0.01, ** *p* < 0.01, *** *p* < 0.001.

Assessment of plasticity kinase signaling in AAV-Hrs-HA-treated animals revealed a 36% increase in phosphorylated PKC substrates (36 ± 11%) and an 11% decrease in PKCα (11 ± 2%) (Figure 7C-D), suggestive of enhanced PKC activity, whereas other kinases (PKCɣ, PKA, and CaMKII) and synaptic proteins (PSD-95, gephyrin, and synaptophysin) were unchanged (Figure 7 and Figure S10). PKC phosphorylates GluA1 at S831, and pGluA1(S831) was markedly elevated by approximately 60% in Hrs-overexpressing animals relative to controls (63 ± 13%), whereas total GluA1 was unchanged (Figure 7). These data suggest that Hrs overexpression selectively enhances PKC activity and GluA1 phosphorylation at S831, thereby enhancing AMPAR conductance.

Collectively, the electrophysiological, proteomic, and biochemical data support Hrs as an important mediator of post-synaptic architecture and glutamatergic function, linking Hrs and ESCRT-0 with the fine-tuning of AMPAR function.

## Discussion

Neuronal ESCRT dysfunction has been linked to neurodegenerative disease, including CHMP2B (ESCRT-III) mutations in patients with frontotemporal dementia (Skibinski et al., 2005) and Hrs (ESCRT-0) in prion disease (Lawrence et al., 2023). Hrs is massively reduced in prion disease, and depleting Hrs from neurons accelerates prion disease progression, including EM-visible structural alterations of the PSD and the build-up of ubiquitinated proteins in synaptic fractions (Lawrence et al., 2023). Yet despite the links between ESCRT proteins and neurodegeneration, little is known regarding how ESCRT-0 spatially and temporally functions at synapses in a non-disease state, or how synaptic activity impacts ESCRT-0 protein expression. Here we show that a major component of ESCRT-0, Hrs, localizes to the pre-synapse as reported, as well as to the PSD. Using biochemical methods and confocal imaging, we observed Hrs as membrane-associated and distributed across the dendritic spine head, neck, and base. Strikingly, Hrs protein expression was dynamically regulated by neuronal activity, increasing with Bic-induced persistent neuronal hyperactivity and decreasing with TTX-induced activity blockade. Furthermore, altering Hrs levels directly impacted AMPAR function and neuronal activity in electrophysiological studies. This is the first evidence that Hrs, and therefore ESCRT-0, functions bidirectionally as a responder and modulator of synaptic activity through facilitating the binding and sorting of ubiquitinated membrane protein cargo into the endolysosomal pathway. While most ESCRT-linked diseases have been investigated at end stage, our data suggest that ESCRT-0 may play an important role in synaptic function and plasticity paradigms prior to observable pathologic states.

As a component of ESCRT-0, Hrs binds and sorts ubiquitinated cargo, thus we evaluated whether disruption of Hrs expression impacts AMPAR trafficking and function, which is known to be regulated by ubiquitination (Schwarz et al., 2010; Goo et al., 2015). We found that stimulation of cultured hippocampal neurons with AMPA led to a rapid co-localization of Nedd4-1 and Hrs within dendrites, suggesting that Hrs is positioned to regulate the trafficking and subsequent degradation of ubiquitinated AMPARs (Schwarz et al., 2010; Scudder et al., 2014; Rodrigues et al., 2016).

Indeed, depleting Hrs from neurons had physiological consequences. Evoked responses in hippocampal slices revealed that Hrs-depleted neurons had a low AMPA/NMDA ratio and a decrease in the AMPAR rise time, consistent with reduced synaptic AMPAR abundance and AMPAR-mediated responses. Our findings suggest that Hrs contributes to the maintenance or delivery of GluA1-containing receptors to the plasma membrane during periods of elevated neuronal activity, providing a potential mechanism underlying the reduced AMPA/NMDA ratio and altered AMPAR kinetics observed following Hrs depletion (Figure. 8).

**Figure 8.**
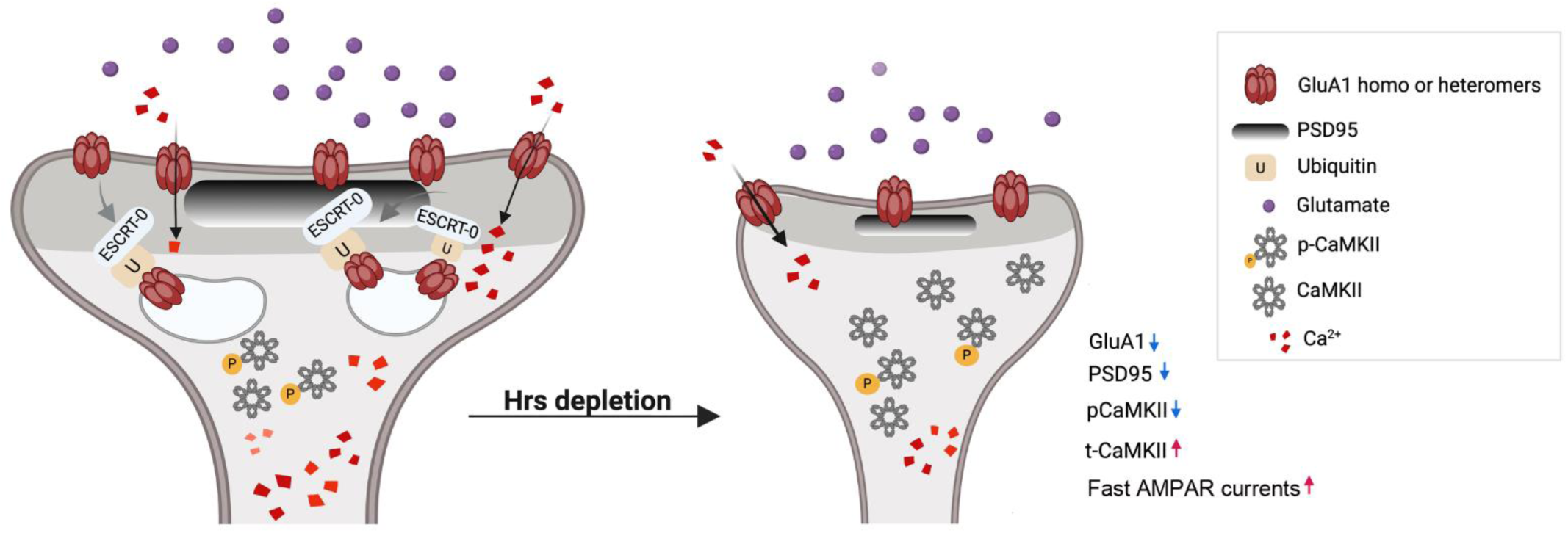
Model of post-synaptic AMPAR regulation by Hrs. Schematic of the post-synapse in control (left) and *Hrs*-deleted (right) neurons shows how ESCRT-0-dependent trafficking may play a role in regulating AMPAR and signaling at the synapse. AMPAR undergo ubiquitination and binding by ESCRT-0 (Hrs/STAM1) for sorting in multivesicular bodies and degradation in lysosomes. *Hrs* depletion was associated with increases in fast AMPAR currents accompanied by reduced phosphorylated CaMKII and PSD-95, suggestive of an LTD-like state.

Phospho-CaMKII-α and –β were markedly reduced (more than 50%) in mice with neuron-specific Hrs deletion (*Hrs^f/f^Syn-Cre*), suggesting that Ca^2+^-dependent signaling in mice may be altered by decreasing Hrs. The low phospho-to-total CaMKII ratio reflects an overall decrease in CaMKII activity. Interestingly, total CaMKII-α and –β levels were elevated by approximately 50%. This dramatic increase could be due to altered turnover of CaMKII. Prior studies showed that active CaMKII-α binds proteasomes (Djakovic et al., 2009; Bingol et al., 2010; Djakovic et al., 2012), and more recently, CaMKII was shown to be ubiquitinated (Quasney and Griffith, 2025), suggesting that CaMKII may be cleared through proteasomes. Given that aberrant proteasomes accumulate during ESCRT dysfunction (Birdsall et al., 2022), depleting Hrs may indirectly reduce proteasomal degradation of ubiquitinated CaMKII, leading to its accumulation. Although increased CaMKII synthesis cannot be excluded, the reduced pCaMKII, PSD95, and gephyrin, together with the electrophysiological data showing a decrease in AMPAR-mediated responses in Hrs-depleted neurons, are consistent with plasticity paradigms that promote a depression of synaptic strength. Evaluating the ubiquitination state and proteasomal turnover of CaMKII in ESCRT-0 dysfunction and disease may further clarify the mechanism underlying the elevated CaMKII.

We previously found that Hrs levels are markedly reduced in the prion-affected brain of humans and experimental mouse models, independent of prion strain (Lawrence et al., 2023). We also found that prion-infected mice show molecular and structural evidence of chronic neuronal hyperactivity, including increased pGluA1-S831 and -S845 and the IEG Arc, as well as the pronounced accumulation of ubiquitinated proteins (Ojeda-Juárez et al., 2022). Prion infection in the Hrs-depleted state accelerates and exacerbates synaptic phenotypes, including a striking expansion of post-synaptic membranes (Lawrence et al., 2023). In light of our current data showing that high neuronal activity leads to an increase in Hrs, the observed reduction in Hrs during prion disease suggests a dysregulated state. Given that ubiquitinated proteins accumulate in other neurodegenerative diseases (Hipp et al., 2019), the mechanisms driving the dysregulated ESCRT and UPS pathways in neurons during disease warrant further investigation.

In conclusion, our studies provide insight into the localization and function of ESCRT-0 in dendrites and at synapses, which is crucial for proper trafficking and turnover of synaptic membrane proteins and receptors. We find evidence supporting a role for post-synaptic ESCRT-0 in modulating synaptic plasticity. In particular, the data show that Hrs expression is dynamically and bidirectionally regulated by neuronal activity. This finding is consistent with the internalization and degradation of synaptic membrane receptors, including AMPARs, by the endo-lysosomal pathway during rapid and long-term neuronal activity. Remarkably, Hrs depletion in neurons alters AMPAR function, and in addition, significantly impacts postsynaptic scaffolds and Ca^2+^-dependent kinases, including CaMKII, suggesting that ESCRT-0 may play a critical role in synaptic plasticity. With abundant evidence for ESCRT dysfunction in neurodegenerative disease, these findings highlight the physiologic contribution of ESCRT-0 in regulating AMPAR function and Ca^2+^-dependent signaling, as well as the dynamic change in Hrs expression during prolonged neuronal hypo- or hyperactivity.

## Materials and Methods

### Animal care

Hrs*^f/f^*mice on a C57BL/6J background (Tamai et al., 2008) were crossed with h*Syn1-Cre* mice (The Jackson Laboratory). The synapsin-1 promoter drives neuron-specific Cre recombinase expression, with robust activity initiating around embryonic day 13 (E13) (Zhu et al., 2001). All *Hrs^f/f^ Cre^+^* and *Cre^−^* littermates were housed under specific pathogen-free conditions in a 12-hour light/dark cycle.

All animal experiments were conducted in strict accordance with protocols designed to minimize animal suffering and were approved by the Institutional Animal Care and Use Committee (IACUC) at the University of California, San Diego (UCSD). Protocols adhered to the guidelines outlined in the *Guide for the Care and Use of Laboratory Animals* issued by the National Institutes of Health.

### Brain collection

Age-matched littermates female *Hrs^f/f^ Cre^+^*and *Cre^−^* mice were euthanized, and brains were immediately removed and dissected along the longitudinal fissure on ice. The right and left cortices were separately snap-frozen in liquid nitrogen and stored at -80 until further use for postsynaptic density (PSD) purification (right) or western blotting (left).

### Post-synaptic density (PSD) fractionation

PSD purification was performed with modifications based on a previously described protocol (Keil et al., 2010) (Fig. S1). Briefly, adult male and female C57BL/6J mice (57-73 days old) were euthanized and fresh cortical and hippocampal tissues were rapidly removed and homogenized (10–15 strokes) on ice using a motor driven glass-teflon dounce-homogenizer in ice-cold HEPES-buffered sucrose solution (0.32 M sucrose, 4 mM HEPES, pH 7.4) supplemented with a 10% (w/v) ratio of cOmplete™ Mini Protease Inhibitor Cocktail (Roche 11836170001) and Pierce™ Phosphatase Inhibitor Mini Tablets (Thermo Scientific™ A32957).

The homogenate was centrifuged at 1000 x g for 15 minutes at 4 °C to remove the nuclear fraction (P1), yielding supernatant S1. S1 was further centrifuged at 10,000 x g for 15 minutes at 4 °C to pellet the crude synaptosomal fraction (P2), and the supernatant (S2) was discarded. The P2 pellet was resuspended in fresh HEPES-buffered sucrose solution and re-centrifuged at 10,000 x g for 15 minutes at 4 °C to yield the washed crude synaptosomal pellet (P2’). P2′ was lysed hypoosmotically by resuspension in 3.6 mL of ultrapure water containing protease and phosphatase inhibitors, followed by gentle homogenization (three strokes, Teflon-glass). The lysate was adjusted to 4 mM HEPES (pH 7.4) and centrifuged at 25,000 x g for 20 minutes at 4 °C (SL50T rotor) to separate the synaptic vesicle-containing supernatant (S3) from the synaptic plasma membrane-enriched pellet (P3). The P3 was then resuspended in 3 mL of ice-cold 50 mM HEPES (pH 7.4) containing 2 mM EDTA and 0.5% Triton X-100, rotated at 4 °C for 15 minutes, and centrifuged at 32,000 x g for 20 minutes at 4 °C (MLS50 swinging-bucket rotor) to yield the PSD-1 fraction. The pellet (PSD-1) was resuspended in 3 mL of the same HEPES-EDTA buffer with 0.5% Triton X-100. A 300 µL aliquot was collected for analysis, and the remaining sample was incubated at 4 °C for 15 minutes and centrifuged at 200,000 x g for 20 minutes at 4 °C, yielding the purified PSD fraction (PSD-2) as the final pellet. The PSD-2 pellet was resuspended in 300 µL of 50 mM HEPES (pH 7.4) containing 2 mM EDTA and 0.5% Triton X-100. Protein concentration was determined using the BCA assay. Equal amounts of protein from each fraction were analyzed by immunoblotting using antibodies against Hrs (Cell Signaling Technology (CST), 15087), PSD95 (CST, 3450), synaptophysin (CST, 5461), and CHMP2B (CST, 76173S).

### Primary neuronal culture

Primary hippocampal or cortical neurons (hippocampal for imaging and cortical for western blot) were prepared from postnatal day one (P1) rat pups using a previously established protocol (Patrick et al., 2003). Briefly, hippocampi or cortical were dissected in ice-cold dissection medium (82 mM Na_2_SO_4_, 30 mM K_2_SO_4_,12 mM MgCl_2_, 20 mM glucose, 0.25 mM CaCl_2_, 1mM HEPES, pH 7.4). Tissues were digested with 18.72 mg papain (Worthington) at 37 °C for 40 minutes. Following enzymatic digestion, cells were gently triturated, centrifuged at 136 x g for 6 minutes, and resuspended in OptiMEM (Gibco) supplemented with 21 mM glucose for plating. Neurons were seeded at a density of 45,000 cells/cm^2^ onto poly-D-lysine (PDL) pre-coated plates (0.05 mg/mL for plastic). After 4 hours, the medium was replaced with growth medium consisting of Neurobasal medium (Gibco) containing 2% B27 (Gibco), 1X L-glutamine (Gibco), and penicillin–streptomycin (1×). Neurons were maintained in a humidified incubator (95% air, 5% CO_2_, 37 °C), and half of the culture medium was replaced every 3–4 days.

To chronically modulate neuronal activity, cortical (western blot) or hippocampal (imaging) neurons at 17 days in vitro (17 DIV) were treated for 72 hours with either 20 µM bicuculline (Bic; Tocris, 0130), 2 µM tetrodotoxin (TTX; Tocris, 1069), or vehicle control (2 µL DMSO).

To assess the effect of shRNA-mediated knockdown on Hrs expression, hippocampal rat neurons were transduced at 5 days in vitro (5 DIV) with 1 µL of shRNA-expressing lentivirus per coverslip. Neurons were maintained for 10 days to allow for effective knockdown and were fixed at 15 DIV for immunocytochemical analysis. Cells were immunostained for Hrs and MAP2 (microtubule-associated protein 2) and processed for fluorescence imaging.

### Immunostaining

Primary hippocampal neurons cultured on coverslips were transduced overnight (16-18h) at 14 DIV with 2 µL of SR2G (attenuated Sindbis virus expressing GFP) for 16 hours to achieve uniform cytoplasmic GFP labeling. At 15 DIV, neurons were rinsed with PBS and fixed for 10 minutes in 4% paraformaldehyde (PFA) at room temperature, followed by three washes in PBS. For co-localization experiment between GFP-Nedd4-1 and endogenous Hrs, neuronal cultures were transduced overnight (16-18h) with SR2G Nedd4-1 (attenuated Sindbis virus expressing GFP-Nedd4-1). Cells were permeabilized and blocked for 20 minutes in 2% bovine serum albumin (BSA) containing 0.25% Triton X-100. After washing, cells were further blocked in 5% BSA for 4 hours at 4 °C. Primary antibodies were applied overnight at 4 °C. Antibodies used included mouse anti-GFP (1:1000, Santa Cruz Biotechnology, sc-9996), anti-Hrs (1:500, Santa Cruz Biotechnology, sc-271455), anti-PSD95 (1:1000, NanoTag Biotechnologies, N3702-Ab635P-L), anti-Bassoon (1:1000, Synaptic Systems 141119), or MAP2 (1:10,000, Abcam, ab5392). Following incubation, coverslips were washed three times in PBS with gentle rocking and incubated for one hour at room temperature with Alexa Fluor–conjugated secondary antibodies (Invitrogen) diluted in 2% BSA. Coverslips were rinsed with PBS and mounted on glass slides using Aqua-Poly/Mount antifade medium (Polysciences). Slides were stored at 4 °C in the dark until imaging.

Neurons were imaged with a Leica DMI6000 B inverted microscope equipped with a VisiTech QLC100 spinning disc confocal scanner and a Ludl 99A041 emission filter wheel. Images were captured using a Hamamatsu ORCA-ER C4742-95 digital camera controlled by a Hamamatsu C4742-95-12ERG camera controller. Excitation was provided by an Andor ILE 03420 laser module integrating 405, 488, 561, and 637 nm laser lines. A Plan Apochromat 63×/1.4 NA oil-immersion objective was used for all imaging. For analysis, Hrs colocalization with PSD95 and Bassoon was assessed within dendritic segments, calculated on a per-cell basis. Colocalization analyses were performed using ImageJ/FiJi and the JaCoP plugin (Bolte and Cordelières, 2006) on maximum-intensity projections using identical acquisition and thresholding parameters across all experimental conditions. For spine-specific analyses, dendritic spines were identified based on morphology, and Hrs puncta were considered spine-localized when overlapping with spine structures. Spine-associated Hrs localization was analyzed separately for PSD95-positive and Bassoon-positive compartments. Furthermore, Hrs fluorescence intensity was quantified in neurons, and normalized to MAP2 signal (binary area) and compared between control and Bic-treated groups.

### Stimulated Emission Depletion (STED) microscopy

3D-STED imaging was used to evaluate Hrs localization with respect to the presynaptic active marker, Bassoon, and postsynaptic excitatory scaffold protein, PSD95. Rat hippocampal neurons were cultured on coverslips (#1.5 thickness, Warner Instruments 64-0732) and transfected at 11 DIV with 1.75 ug hSyn1-Lck-GFP (Addgene #177333) using the CalPhos Mammalian Transfection Kit (Clonetech, 631312). After 48 hours, neurons were fixed with 4% (v/v) PFA (Electron Microscopy Sciences) for 20 minutes at room temperature and washed three times with 0.1 M PBS. Coverslips were permeabilized for 20 minutes in a 0.25% Triton-X (Fisher) in PBS solution and washed three times with 0.1 M PBS. The samples were blocked with 5% w/v BSA in 0.1 M PBS for four hours. After blocking, coverslips were washed with PBS and incubated overnight at 4 °C with primary antibodies diluted in 2% BSA in 0.1 M PBS. The following primary antibodies were used: Anti-GFP Atto488 (NanoTag, N0304-AT488-L,1:250), anti-Bassoon (chicken; Synaptic Systems, 141119,1:1000), anti-PSD95 Abberior STAR635P (NanoTag, N3702-Ab635P-L, 1:1000) and anti-Hrs (Mouse, SantaCruz sc-271455, 1:500). The next day, coverslips were washed three times with 0.1 M PBS and incubated for one hour at room temperature with secondary antibodies diluted in 2% BSA in 0.1 M PBS. Secondary antibodies were goat anti-chicken STAR460L (Abberior, 1005-500UG, 1:200) and goat anti-mouse Alexa 594 (Thermo Fisher, A-11005, 1:200). Coverslips were washed in 0.1 M PBS three times and mounted with Fluoromount-G (SouthernBiotech, 0100-01). Three independent primary neuronal cultures were prepared, with 10 neuronal dendrites imaged per culture.

3D-STED imaging was conducted on the Abberior Facility Line microscope (Abberior Instruments GmbH, Germany), which operates in both confocal and STED modalities. Image acquisition was performed with Abberior Lightbox software. For confocal acquisition, anti-GFP Atto 488 was excited at 488 nm (detection range: 498 – 551 nm), selecting neurons with lower levels of GFP to minimize spectral bleed-through with STAR 460L when imaged in conjunction. For STED acquisition, STAR 460L was excited with a pulsed source at 440 nm (detection range: 580 - 630 nm), AF594 was excited at 561 nm (detection range: 580 - 630 nm), and STAR635P was excited at 640 nm (detection range: 650 - 755 nm). All STED channels were deactivated with a pulsed source at 775 nm (10% deactivation power) and underwent TIMEBOW (fluorescence lifetime imaging) to capture lifetime information for post-processing. A pixel dwell time of 5 μs was employed, and the pinhole was set to 1.3 Airy Units at 574 nm. An oil-immersion objective lens (UPLXAPO 60X oil immersion lens, 1.42 NA, Olympus, Japan) was used for all STED imaging. All images were deconvolved in Huygens Essential software prior to analysis in Aivia. A representative image was used to train Pixel Classifiers and generate 3D Object Segmentation Recipes for PSD-95, Bassoon, and Hrs, which were then applied to all images using batch processing. Spatial relationship analyses were subsequently performed using nearest-neighbor measurements within a 0.3 μm search radius, including centroid-to-centroid and shell-to-shell distance calculations.

### Surface biotinylation assay

Primary cortical neurons were prepared from Hrs^f/f^ mice and cultured. At DIV 7, neurons were transduced with either AAV-Cre [pENN.AAV.hSyn.HI.eGFP-Cre.WPRE.SV40 (AAV PHP.eB), Addgene #105540-PHPeB] to induce Hrs deletion or control AAV-YFP [hSyn1-eYFP (AAV PHP.eB), Addgene #117382-PHPeB]. Following 6–10 days of viral expression, neurons were treated with Bic (20 μM) or DMSO vehicle control for 24 hours.

Cell-surface proteins were labeled using the membrane-impermeable reagent Sulfo-NHS-SS-Biotin (thermos #21335). Briefly, neurons were gently washed with progressively cooled PBS containing 1 mM MgCl₂ and 2.5 mM CaCl₂ (PBS^++^) and maintained at 4°C throughout the labeling procedure to prevent membrane trafficking and receptor internalization. Neurons were incubated with NHS-LC-Biotin (1 mg/mL in PBS^++^) for 15 minutes at 4°C. Following biotinylation, cells were washed twice with ice-cold PBS^++^ containing 0.1% bovine serum albumin (BSA) to remove excess reagent. For protein extraction, neurons were lysed in precipitation buffer containing 100 mM NaCl, 10 mM Na₂HPO₄, 5 mM EDTA, 5 mM EGTA, 1% Triton X-100, 0.1% SDS, protease inhibitors (Thermo Fisher Scientific), and PhosSTOP phosphatase inhibitors (Roche). Lysates were rotated at 4°C for 20 minutes and clarified by centrifugation at 14,000 × g for 20 minutes. An aliquot (10–20%) of each lysate was retained as the total protein input fraction. Protein concentrations were determined by bicinchoninic acid (BCA) assay and lysates were normalized to equal protein concentrations. Surface-biotinylated proteins were isolated by incubation with NeutrAvidin beads (Pierce #29200, 100 μL per sample) overnight at 4°C with rotation. Beads were pre-washed three times in lysis buffer containing 1% Triton X-100 before use. Following incubation, beads were washed three times with lysis buffer and bound proteins were eluted by boiling in 3× SDS sample buffer at 95°C for 10 minutes. Eluted surface fractions and corresponding input lysates were immunoblotted with antibody against GluA1.

### Electrophysiology

Organotypic hippocampal slices were prepared from C57BL/6 mice as described previously (Dore et al., 2021). Slices were injected at 2-3 DIV with either a scrambled control shRNA AAV or an Hrs-targeting shRNA AAV (VectorBuilder, Chicago, IL) to knockdown Hrs expression. Slices were used for electrophysiological recordings at 7-14 DIV [5-12 days post-infection (DPI)]. AMPA/NMDA ratios in slices recorded at 5–8 DPI and 9–12 DPI showed efficient Hrs knockdown.

A surgical cut on the CA3 region was done on organotypic slices to prevent stimulus-induced bursting. Slices were transferred into the recording chamber with a continuous flow of oxygenated artificial cerebrospinal fluid (aCSF), containing 119 mM NaCl, 2.5 mM KCl, 26 mM NaHCO_3_, 1 mM NaH_2_PO_4_, 10 mM glucose, 4 mM CaCl_2_, 4 mM MgCl_2_, 10 μM gabazine, and 4 μM 2-chloroadenosine (pH 7.4). The bath aCSF was perfused at a rate of 1.5-2.0 ml/min and gassed with 5% CO_2_/95% O_2_ at 28-32 °C. The slices were allowed to rest in the chamber for 5-10 min before all recordings. CA1 pyramidal neuron layer was identified under the microscope of the recording rig. The stimulating electrode (contact Pt/Ir cluster bipolar electrodes (Frederick Haer) was placed in Stratum Radiatum about 300 μm down the apical dendrite of CA1 pyramidal neurons. GFP-positive neurons were selected from the pyramidal neuron layer in CA1 for patch clamp recording. Borosilicate glass pipettes (outer diameter: 1.5 mm, Warner, Hamden, CT) were pulled from a micropipette puller (Model P-97, Sutter). The recording electrodes had a resistance of 2-5 MΩ when filled with an internal solution containing 115 mM Cesium methanesulfonate, 20 mM CsCl, 10 mM HEPES, 2.5 mM MgCl_2_, 4 mM Na_2_ATP, 0.4 mM Na_3_GTP, 10 mM sodium phosphocreatine (Sigma), and 0.6 mM EGTA (Amresco), at pH 7.25. Cesium-based internal solutions were used to block potassium conductances and optimize space clamp in voltage-clamp whole cell recordings (Fourgeaud et al., 2010) and were applied consistently across all experimental groups. A MultiClamp 700B amplifier, an Axon Digidata 1550B, and Clampex 11 software (Molecular Devices, San Jose, CA, USA) were used for data acquisition, digitized at 2-10 kHz, and filtered at 2 kHz. The liquid junction potential was compensated for prior to forming the cell-attached mode for all recordings. A minimum of 2 GΩ seal resistance was required before rupturing the membrane for whole-cell configuration. For each slice, the input-output test was done first to define the stimulation intensity to evoke AMPAR or NMDAR responses. 50∼70% of the stimulation intensity inducing the maximum AMPAR response was used for excitatory post-synaptic currents (EPSCs) in the same slice. Evoked AMPAR or NMDAR-mediated EPSCs were recorded under voltage clamp at a holding potential of −60 mV or +40 mV, respectively. The series resistance was tested by “membrane test” in Clampex 11, and the recordings with series resistance larger than 30 MΩ or with variations larger than 30% were excluded from data analysis (Fourgeaud et al., 2010). These inclusion criteria were applied uniformly across experimental conditions.

Evoked AMPAR or NMDAR EPSCs of each recorded cell were averaged from 30–50 recorded sweeps, excluding sweeps with bursting or excessive noise. The amplitudes of AMPAR or NMDAR EPSCs, rise/decay time, and slope of AMPAR EPSCs were analyzed using Clampfit 11 software (Molecular Devices, San Jose, CA, USA). AMPAR currents were classified as ‘fast’ if the majority of the sweeps (more than 90%) had one peak and decayed within 20 ms from the start of the sweep. In contrast, if more than 10% of the sweeps showed multiple peaks and had decay times greater than 25 ms, the recording was defined as a slow current. After this classification, sweeps recorded for each neuron were averaged, and the peak amplitude, rise and decay time, and slope were calculated from the averaged current. We note that minimal variability in the latency of evoked currents was observed across recordings.The data with averaged NMDAR EPSCs smaller than 5 pA were excluded from analysis. AMPA/NMDA ratio was calculated using the absolute value of the peak AMPAR EPSC amplitude divided by NMDAR EPSC amplitude measured at 100 ms after stimulation (Fourgeaud et al., 2010). The kinetics of the EPSCs from each recorded neuron were analyzed in Clampfit–Statistics, measuring 20–80% rise time/slope and 100–37% decay time/slope.

### Mass spectrometry-based proteomics analysis

Post-synaptic membrane (PSM) fractions were prepared with modifications to a previously described protocol (Thakar et al., 2017). All steps were performed at 4 °C using pre-chilled buffers supplemented with protease (cOmplete Mini, Roche) and phosphatase (PhosSTOP, Roche) inhibitors, which were added fresh before use. Cortical tissue from Hrs^f/f^Syn-Cre^+^ and Cre^-^ mice was homogenized to a 10% (w/v) suspension in Buffer A (0.32 M sucrose, 1 mM MgCl₂, 0.5 mM CaCl₂, 1 mM NaHCO₃) using a glass Dounce homogenizer, and the homogenates were centrifuged at 700 × g for 30 minutes to remove nuclei and debris. The resulting supernatant was centrifuged at 13,800 × g for 10 minutes to yield the crude synaptosome pellet (P2), which was resuspended in Buffer B (0.32 M sucrose, 1 mM NaHCO₃) and loaded onto a discontinuous sucrose gradient consisting of 1.4 M sucrose (Buffer D, bottom), 1.0 M sucrose (Buffer C, middle), and Buffer B (top). Gradients were centrifuged at 82,000 × g for 1 hour, and synaptosomes were collected from the cloudy band at the 1.0/1.4 M sucrose interface. The fraction was subjected to hypoosmotic shock in Buffer E (1 mM NaHCO₃, 0.5× inhibitors) and incubated on ice for 10 minutes, followed by solubilization in Buffer F (0.32 M sucrose, 1% Triton X-100) for 15 minutes at 4 °C. Samples were then centrifuged at 82,000 × g for 1 hour, and the resulting pellets corresponding to PSMs were resuspended in 50–100 µL of Buffer F and stored at −80 °C.

In preparation for mass spectrometry analysis, isolated synaptosome protein (74 μg) from cortical tissue was precipitated with methanol and chloroform. The precipitated proteins were digested with trypsin following a previously described protocol (Kawata et al., 2023). The digested peptides were desalted and dried using a Speed-Vac. Each peptide sample was labeled with Thermo Scientific TMT 10plex isobaric tags according to an established method (Zecha et al., 2019). After labeling, the TMT-labeled samples were combined into a single tube, with 100 μg set aside for unmodified peptide analysis, while the remaining portion was used for phosphopeptide enrichment. Phosphorylated peptides were enriched sequentially using ferric nitrilotriacetate (Fe-NTA) Thermo Scientific Phosphorylation Enrichment Kit. Both the TMT-labeled unmodified and phosphorylated peptides were fractionated offline using high-pH reverse-phase spin columns (Thermo Scientific).

The TMT labeled samples were analyzed on an Orbitrap Fusion Lumos Tribrid Mass Spectrometer (Thermo Scientific). Peptides were injected onto a 25 cm, 100 μm ID column packed with BEH 1.7 μm C18 resin (Waters) and separated at a flow rate of 300 nL/min using an EasynLC 1200 system (Thermo Scientific). The mobile phase consisted of Buffer A (0.1% formic acid in water) and Buffer B (90% acetonitrile in 0.1% formic acid). The gradient program was as follows: –25% B over 120 min, an increase to 40% B over 40 min, an increase to 100% B over 10 min, followed by a 10 min hold at 100% B, for a total run time of 180 min.

Peptides were eluted directly from the column tip and nanosprayed into the mass spectrometer using a 2.5 kV voltage applied at the column’s back end. The Orbitrap Fusion Lumos was operated in data-dependent acquisition (DDA) mode. Full MS1 scans were acquired in the Orbitrap at 120,000 resolution. The cycle time was set to 3 seconds, during which the most abundant precursor ions per scan were selected for collision-induced dissociation (CID) MS/MS in the ion trap. MS3 analysis with multi-notch isolation (SPS3) was used to detect TMT reporter ions at 60,000 resolution. Monoisotopic precursor selection was enabled, and dynamic exclusion was set to 10 seconds to minimize repeated sampling of abundant ions.

Mass spectrometry data were analyzed using Proteome Discoverer 2.5 to identify peptides and phosphorylation sites. The MS/MS spectra were searched against the Uniprot mouse protein database with isoforms (version 2022-06-14), along with a common contaminant protein list. A decoy database (reverse-sequence Uniprot database) was used to filter identifications to a 1% false discovery rate (FDR).

Peptides were allowed a maximum of two miscleavages and semi-tryptic cleavage sites. The search included TMT labeling as a static modification on lysine and peptide N-termini (+229.162932 Da) and carbamidomethylation of cysteine (+57.021464 Da). Phosphorylation (+79.9663 Da) was included as a variable modification on serine, threonine, and tyrosine. TMT reporter ion correction factors were applied based on the specific lot number of the TMT reagent. For unmodified peptide quantification, the average reporter S/N threshold was set to 10, and the SPS mass matches [%] threshold was set to 65. For phosphopeptide quantification, the SPS mass matches [%] threshold was set to zero, and a signal-to-noise (S/N) threshold of 100 was applied as previously described. The phosphorylation site localization threshold was set at 75%.

### Mass spectrometry data processing and visualization

Data analysis was performed in R [version 4.4.2 (2024-10-31)]. Unmodified proteome data was imported into R as tandem mass tag intensities associated with gene symbols and UniProt identifiers. Phosphopeptide data was imported into R as tandem mass tag intensities associated with gene symbols, UniProt identifiers, peptide sequence information, and false discovery rate (FDR) confidence scores. Both high and low confidence FDR phospho-peptides were considered. Only a single uniprot ID was considered for phosphopeptides associated with more than one uniprot ID. Phosphopeptide intensities were normalized to unmodified protein intensities within each sample. Phosphopeptides assigned to proteins not detected in the unmodified proteome data were excluded. Fold-changes and log2 fold-changes for unmodified proteins and phosphopeptides were calculated by dividing the mean intensity of Cre+ samples by the mean intensity of Cre- samples. Statistical significance of differences was assessed using two-sample student’s t-test with Welch’s approximation to the degrees freedom. Abundance differences were considered significant if the unadjusted p-value was less than 0.05. Heatmap clustering was performed in R-package pheatmap (version 1.0.13) using a single linkage method. Gene ontology (GO) and Kyoto Encyclopedia of Genes and Genomes (KEGG) pathway were performed on significantly different proteins or phosphopeptides using the R-package clusterProfiler (version 4.12.6) using all genes listed in the Org.Mm.eg.db package database (version 3.19.1) as background. Only uniprot identifiers that could be mapped to entrez IDs by the *bitr()* command in clusterProfiler were considered in GO and KEGG analysis.

Mass spectrometry data was visualized in R (version 4.5.2 (2025-10-31)) using the R-packages ggplot2 (version 4.0.1), ggrepel (version 0.9.6), pheatmap (version 1.0.13), patchwork (version 1.3.2), and scales (version 1.4.0) with additional formatting performed in Inkscape (https://www.inkscape.org, version 1.4).

Raw mass spectrometry data is publicly available in the ProteomExchange depository (PXD073715).

### AAV production

AAVs were generated according to established protocols (Challis et al., 2019). Briefly, HEK293T cells (ATCC) were triple transfected with capsid, viral genome, and helper plasmids using polyethylenimine. Five days after transfection, virus from both media and cell lysates were collected and purified via ultracentrifugation in an iodixanol (OptiPrep, Sigma Aldrich) density gradient. Viral titer was determined by qPCR against a standard curve using primers that anneal to the viral genome’s WPRE element.

### AAV transduction

CAP-B10 hSyn-GFP and CAP-B10 hSyn-Hrs viral vectors were transduced in male C57BL/6J mice (80–120 days old) by administering 1.11 × 10¹² vg by intravenous retro-orbital injection into the venous sinus. Mice were euthanized 21 days post-transduction, and cortical tissues were collected for biochemical analyses.

### Immunoblot analysis

The cortical tissues were homogenized in ice-cold RIPA lysis buffer (50 mM Tris-HCl pH 7.4, 150 mM NaCl, 1 mM EDTA, 1% NP40, 0.5% DOC, 0.1% SDS) supplemented with 2% N-lauryl sarcosine, containing protease and phosphatase inhibitors, and Benzonase^TM^. A 10% weight-to-volume (w/v) lysate was generated using a beadbeater (BioSpec). Lysates were incubated on ice for 30 minutes, followed by centrifugation at 7000 x g for 10 minutes. Supernatants were collected, and protein concentrations were determined using a BCA assay. For western blot analysis, samples (DTT-reduced) were analyzed using the following antibodies: anti-Hrs (CST, 15087), pGluA1-S845 (CST, 8084), pGluA1-S831 (CST, 75574), GluA1 (CST, 8084), stargazin (CST, 8511), pGluA2-S880 (Phosphosolutions, p1170-880), GluA2 (CST, 13607), pGluN2B-S1303 (CST, 71335), pGluN2B-Y1472 (CST, 4208), GluN2B (CST, 14544), GluN2A (CST, 4205), p(Ser) PKC substrates (CST, 2261), PKC-α (CST, 2056), PKC-γ (Santa Cruz, SC-211), PKA (CST, 9621), pCaMKII-T286 (CST, 12716), CaMKII (CST, 4436), PP2B (CST, 2614S), gephyrin (CST, 14304), PSD95 (CST, 3450), synaptophysin (CST, 36406), ubiquitin (CST, 3936), alpha-tubulin (CST, 3873), and GAPDH (Genetex, GTX627408-01). Signals were detected using a chemiluminescent substrate, captured and quantified using the Fuji LAS 4000 imager, and quantified using Multigauge V3.0 software.

### Data analysis, statistics, and reproducibility

Quantitative data were analyzed using GraphPad Prism (version 10). Data distribution was assessed for normality. For comparisons between two groups, unpaired two-tailed Student’s *t* tests were used for normally distributed data. Data are presented as mean ± SEM. All imaging analyses were performed blind to experimental condition, and experiments were replicated across multiple independent cultures or animals to ensure reproducibility.

For biochemical fractionation experiments, protein levels were compared using unpaired two-tailed Student’s *t* tests. For immunocytochemistry analyses, colocalization of Hrs with PSD95 or Bassoon was quantified using Pearson’s correlation coefficient, and group differences were assessed using unpaired two-tailed *t* tests. Hrs localization within dendritic spines was quantified as (i) the percentage of spines containing Hrs and (ii) the distribution of Hrs puncta across spine head, neck, and base compartments; statistical comparisons was performed using one-way ANOVA with Tukey’s multiple comparison test for compartmental distribution analyses of Hrs in spines. For GFP–Nedd4-1 experiments, colocalization between Hrs and GFP–Nedd4-1 was quantified using Manders’ overlap coefficients with thresholding, reporting both the fraction of Hrs overlapping with GFP–Nedd4-1 and the fraction of GFP–Nedd4-1 overlapping with Hrs; comparisons between control and AMPA-treated conditions were performed using unpaired two-tailed *t* tests.

For Western blot analyses of primary neurons treated with TTX or bicuculline, isolated synaptosomes, Hrs^f/f^Syn Cre^+^ and Cre^-^, and AAV-Hrs-HA and AAV-GFP injected animals, protein levels were quantified by densitometry, normalized to loading controls, and compared using unpaired two-tailed Student’s *t* tests. Electrophysiological data was obtained from hippocampal organotypic slices expressing either scrambled shRNA or Hrs shRNA. AMPA/NMDA ratio, normalized AMPAR-mediated current amplitude, AMPAR rise time, decay time, and decay slope were quantified and compared between groups using unpaired two-tailed Student’s *t* tests. The percentage of cells with slow AMPAR EPSCs was compared using the Chi-square (Fisher’s exact) test. Cells were considered independent experimental units. Recordings were obtained from five separate cultures.

Phosphoproteomics data were analyzed using Gene Ontology (GO) enrichment analysis, volcano plots to visualize differentially phosphorylated proteins, and heatmap-based hierarchical clustering to assess global phosphorylation patterns across conditions.

## Data availability

All data generated or analyzed during this study are included in the manuscript and supporting files.

## Acknowledgments

We thank the animal care staff at the University of California, San Diego, for excellent animal care.

## Author contributions

MP, LD, GNP, and CJS conceptualized and designed the experiments; MP, LD, AW, SG, JER, KS, and DPP performed biochemistry, immunolabeling, and/or confocal microscopy experiments. DBM performed the mass spectrometry experiments under the guidance of JRY. YD and KD performed and analyzed the electrophysiology experiments. MKBM and JHT performed the STED imaging experiments. EES and TFS generated the AAV viruses, and BA transduced the mice, under the guidance of VG and SR, respectively. JW performed and supervised all mouse management and genotyping. MP, LD, DBM, AW, YD, JEM, SG, JER., JHT, KD, MKBM, GNP, and CJS analyzed data. The manuscript was written by MP, GNP, and CJS, with input from all authors.

## Supplementary Figures

**Figure S1.**
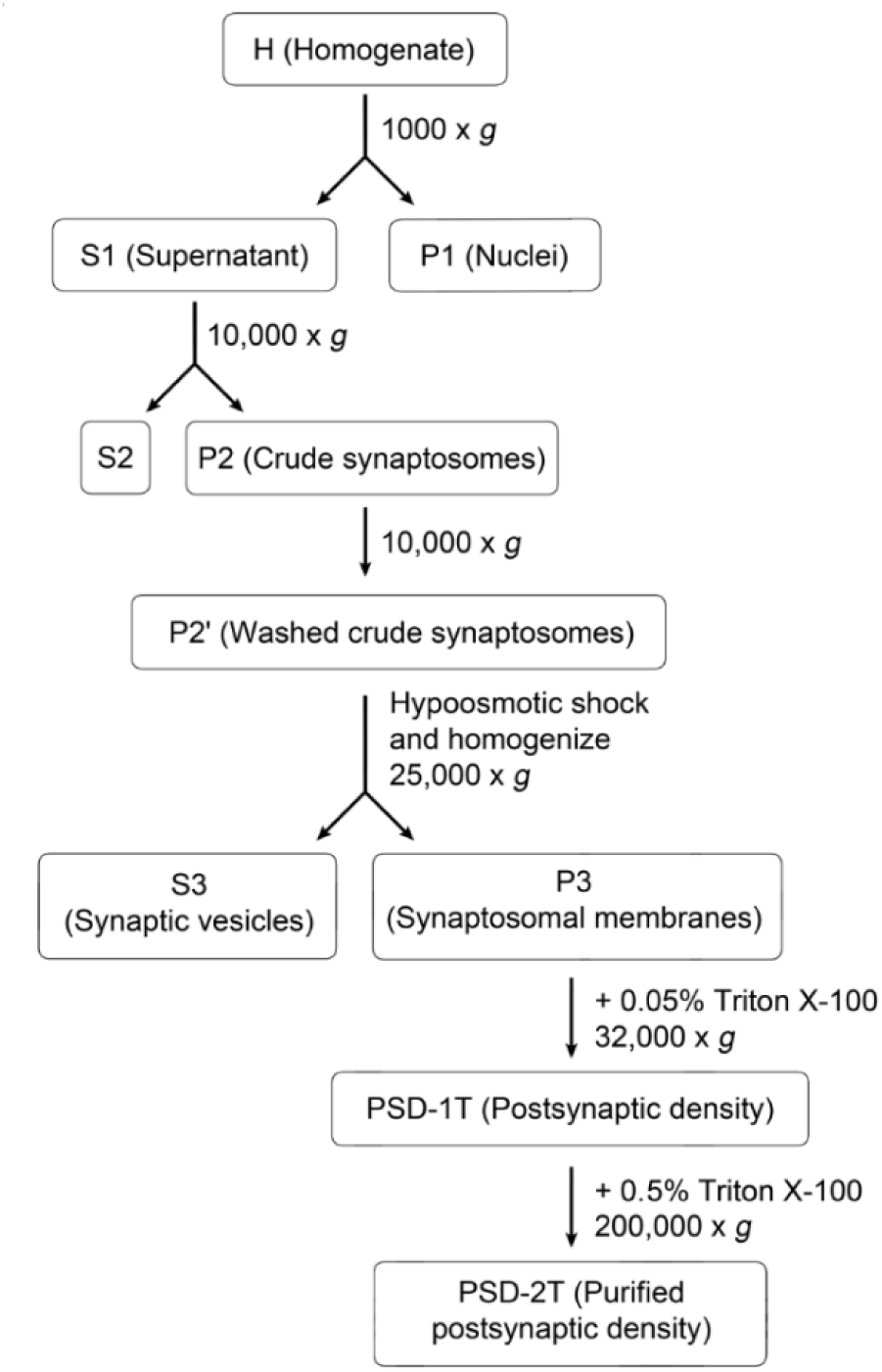
Schematic diagram depicting post-synaptic membrane (PSD2) purification protocol from WT mouse brain (cortex and hippocampus).

**Figure S2.**
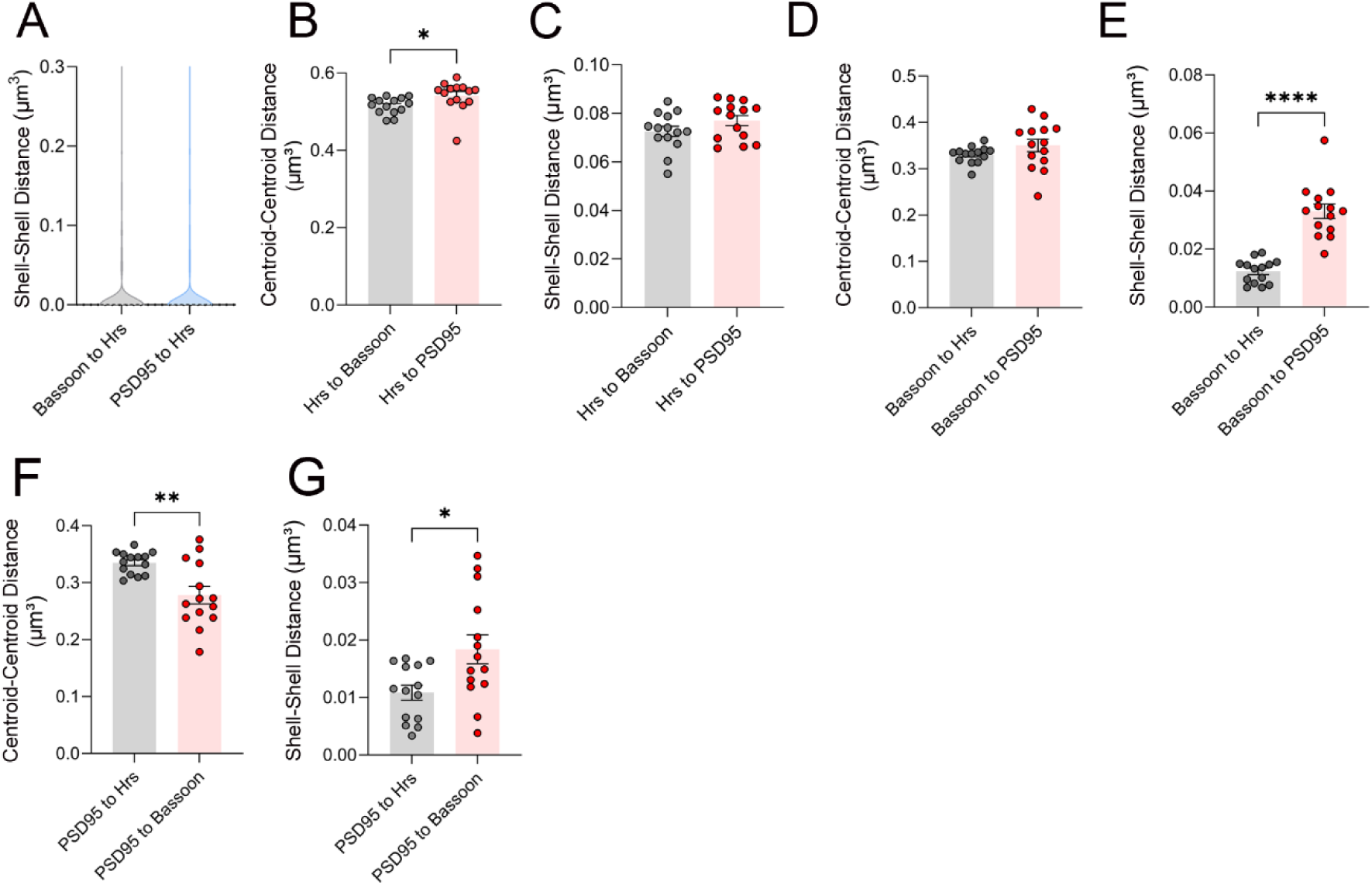
Nanoscale spatial relationship of Hrs with synaptic markers. (A–G) Quantification of nanoscale distances between Hrs and synaptic markers using super-resolution imaging. (A, C, E, G) Shell-to-shell distance measurements represent the minimum edge-to-edge distance between segmented objects. (B, D, F) Centroid-to-centroid measurements represent the distance between object centers. Distances were calculated between Hrs and the presynaptic marker Bassoon and between Hrs and the postsynaptic marker PSD-95. Additional pairwise analyses comparing Bassoon and PSD-95 are shown. Data are presented as mean ± SEM. Statistical significance was determined using two-tailed unpaired t-tests (\**p* < 0.05, \*\**p* < 0.01, \*\*\**p* < 0.001).

**Figure S3.**
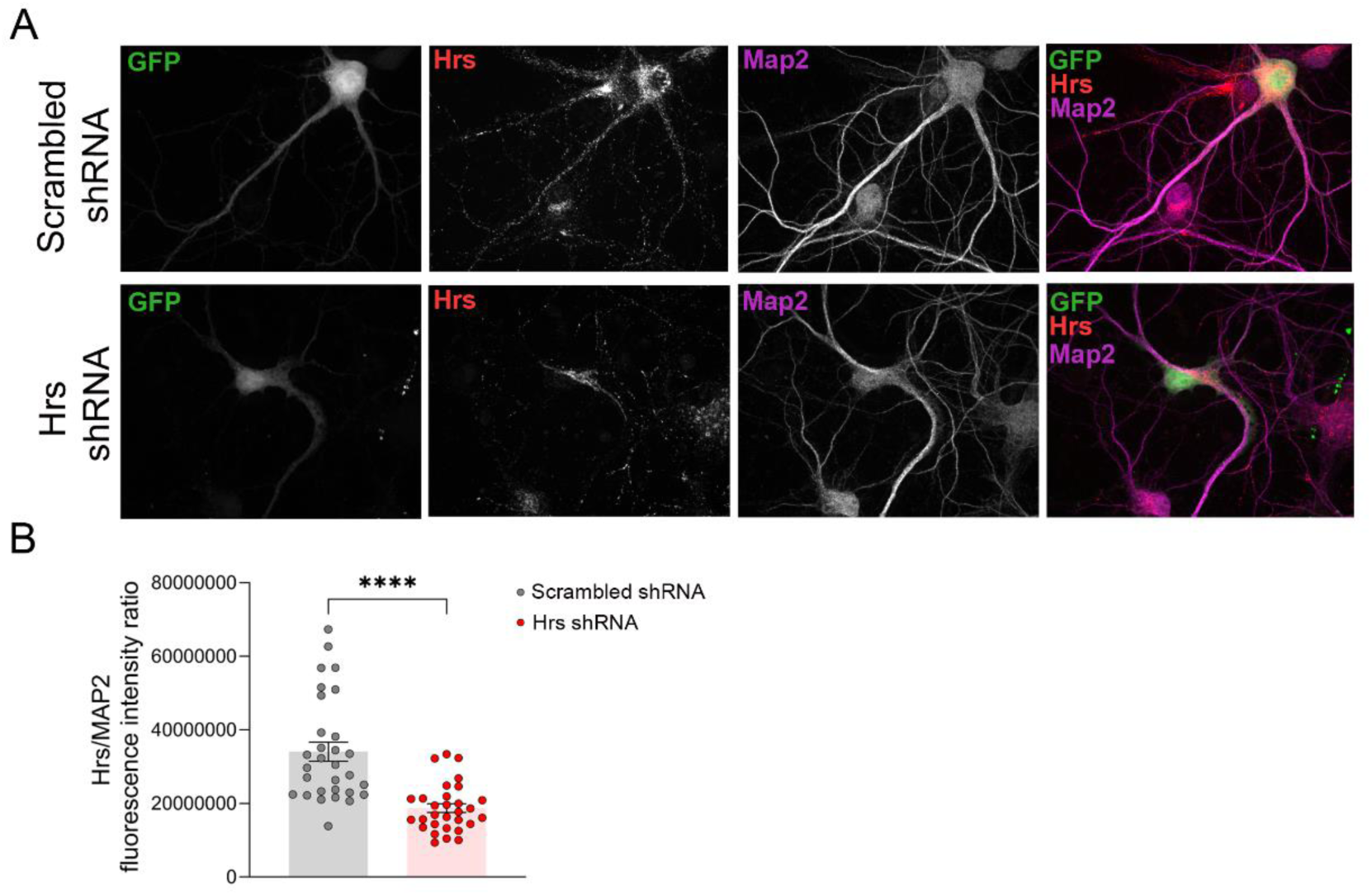
shRNA-mediated knockdown and immunostaining of Hrs in primary hippocampal neurons. Primary hippocampal neurons were transduced with shRNA-expressing lentivirus (Hrs-shRNA or scrambled shRNA) at 5 days in vitro (5 DIV) using 1 µL of virus per coverslip. At 15 DIV, neurons were fixed and immunostained for Hrs, MAP2, and GFP. Hrs/Map2 fluorescence intensity was quantified, revealing a significant reduction in Hrs levels in the Hrs-shRNA–treated neurons compared with control neurons. Unpaired two-tailed *t*-test. **** *p* < 0.0001.

**Figure S4.**
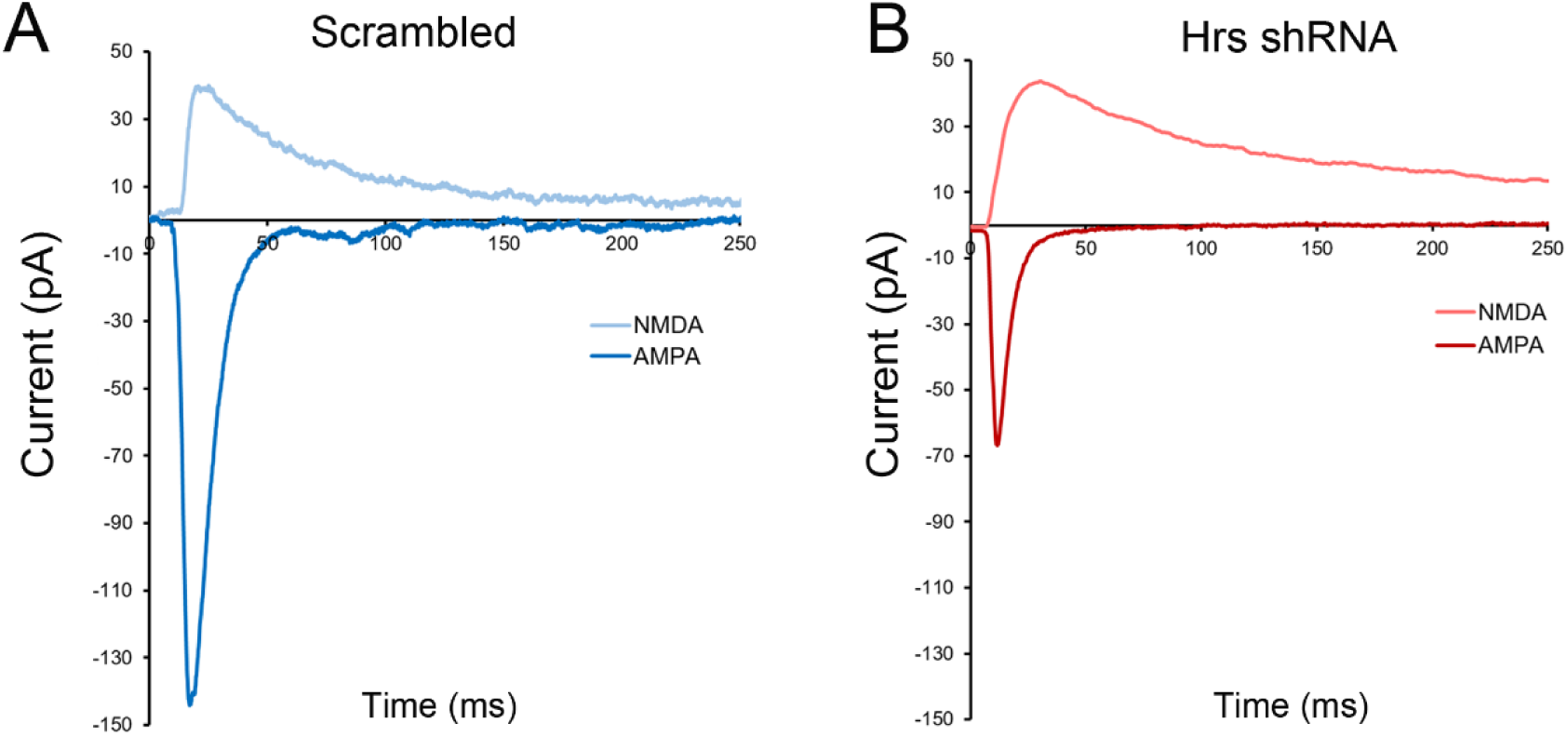
Example traces of NMDA and AMPA currents from the same neuron.

**Figure S5.**
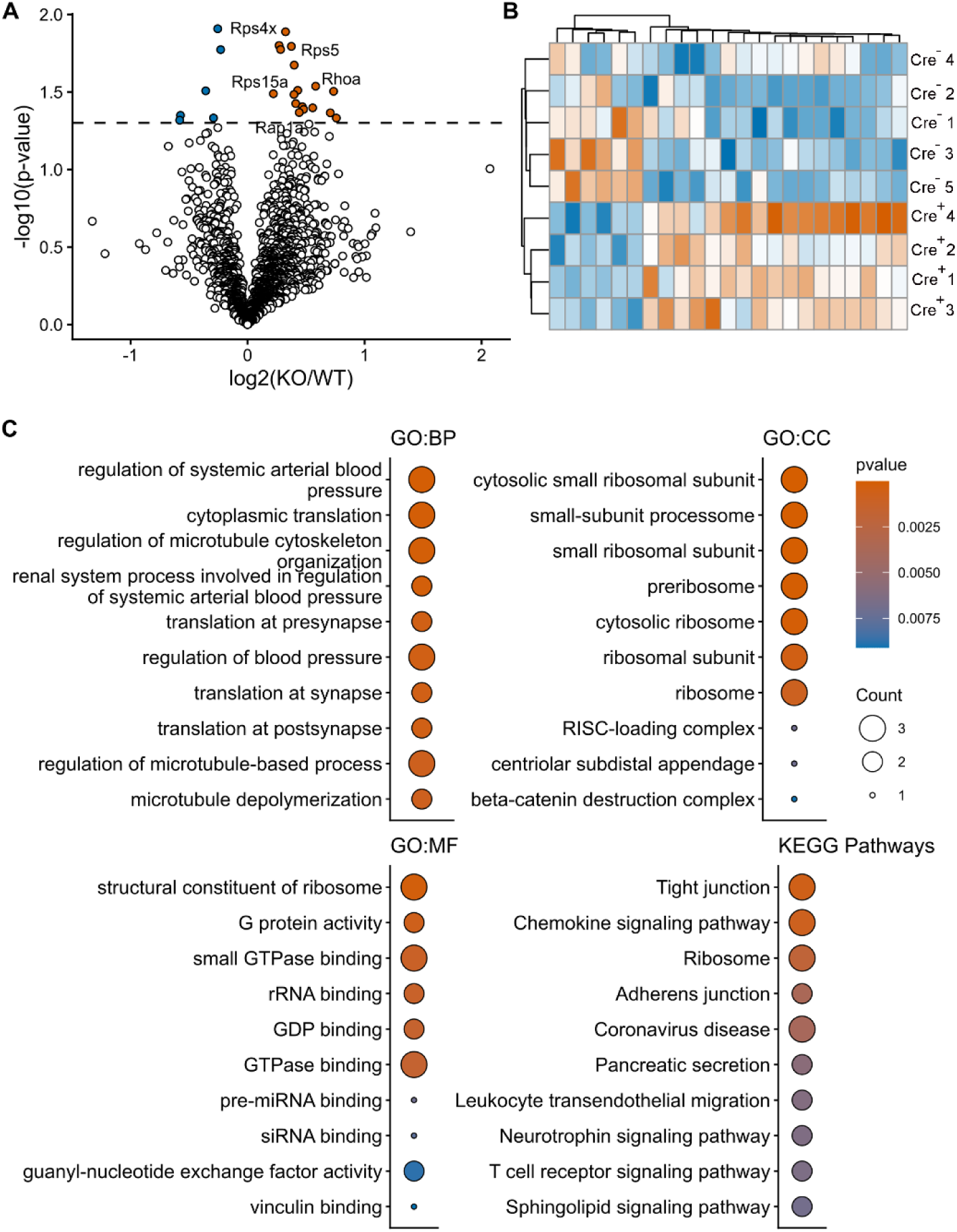
Differential protein abundance and functional enrichment in the unmodified proteome of Hrs-depleted post-synaptic membranes from mouse forebrain (cortex and hippocampus). (A) Volcano plot showing differentially abundant proteins in *Hrs^f/f^ Syn1-Cre^+/-^* and *Hrs^f/f^Syn1-Cre^-/-^*post-synaptic density (PSD2 fraction). The dashed horizontal line indicates the significance threshold at –log10 (0.05). A total of 2,064 proteins were identified (including splice variants), with 23 showing significant changes (17 increased and 6 decreased; *p* < 0.05). (B) Heatmap representing relative abundance of differentially expressed proteins across samples, hierarchically clustered. Color gradient indicates intensity of scaled relative peptide abundance (vermilion = high, blue = low). Sample labels correspond to clustering order (Cre^-^1 to Cre^-^5 and Cre^+^1 to Cre^+^4). (C) Gene ontology (GO) classified gene sets show top biological process (BP), cellular component (CC), and molecular function (MF). KEGG enrichment analyses are also shown. The color gradient indicates unadjusted *p*-values. Point size corresponds to the number of genes enriched within each ontology or pathway.

**Figure S6.**
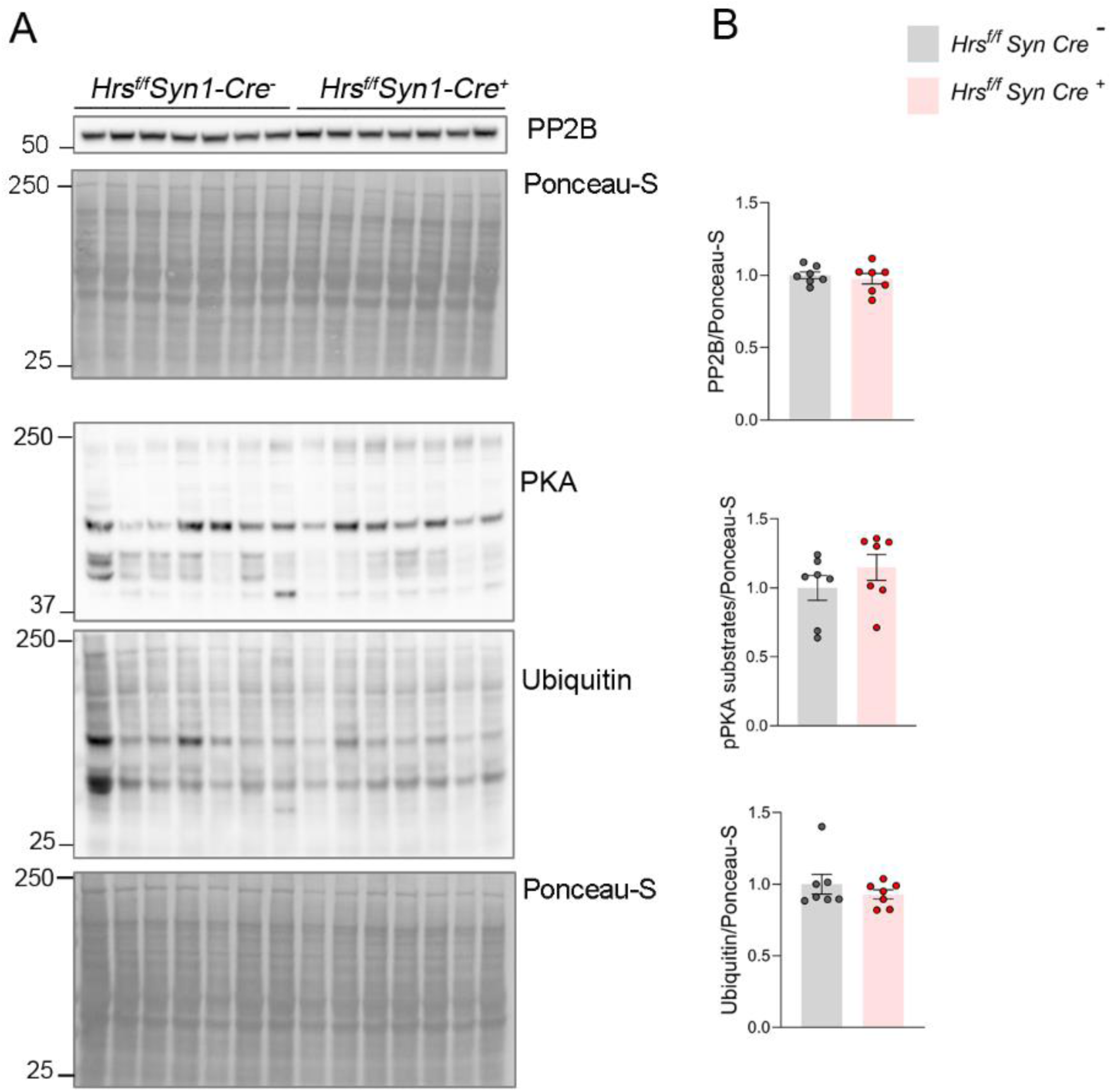
Neuronal Hrs depletion in mice did not impact the phosphorylated PKA substrate, PP2B, or ubiquitin levels. (A) Western blots of cortical lysates from *Hrs^f/f^Syn1-Cre^-/-^*and *Hrs^f/f^Syn1-Cre^+/-^*mice for PKA substrates, PP2B, and ubiquitin (n = 7 per group). (B) Densitometric quantification of western blot signals. Mean ± SEM. Unpaired two-tailed *t*-test.

**Figure S7.**
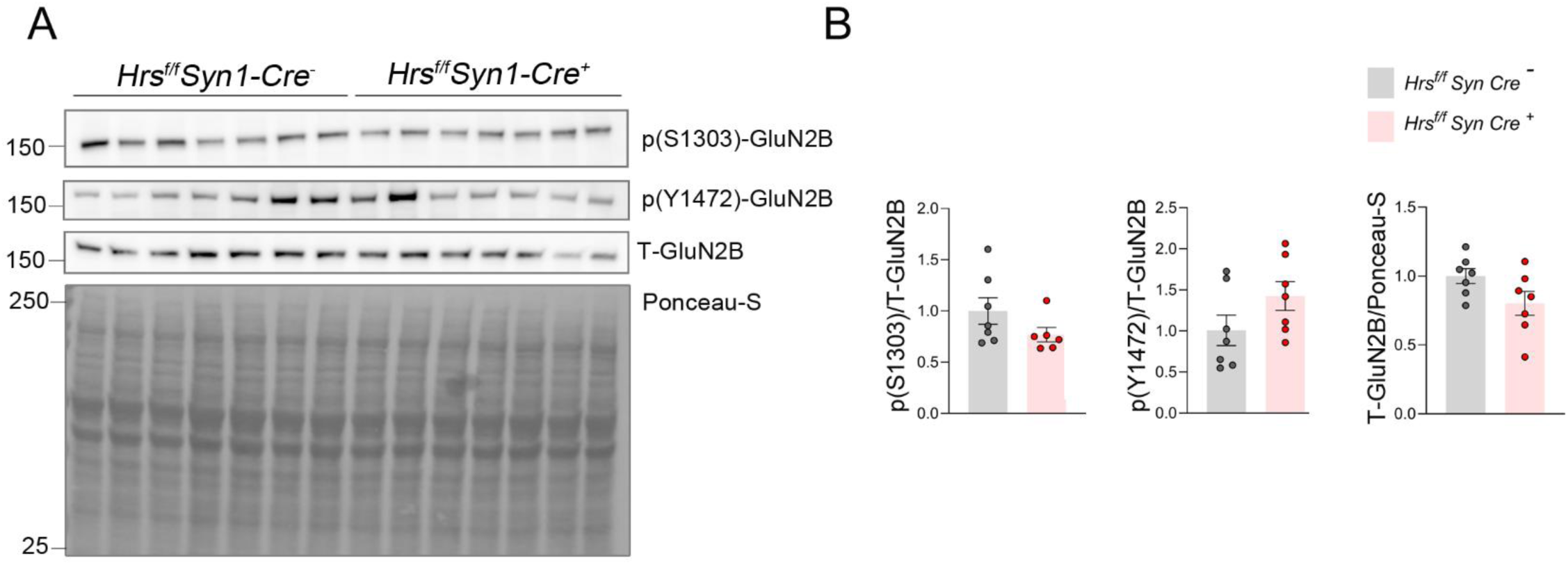
Neuronal Hrs depletion did not change the phosphorylation status of NMDA receptors. (A) Western blots of cortical lysates from *Hrs^f/f^Syn1-Cre^-/-^*and *Hrs^f/f^Syn1-Cre^+/-^* mice for GluN2B receptor and phosphorylation at S1303 and Y1472 (n = 7 per group). (B) Densitometric quantification of western blot signals. Mean ± SEM. Unpaired two-tailed Student’s *t*-test.

**Figure S8.**
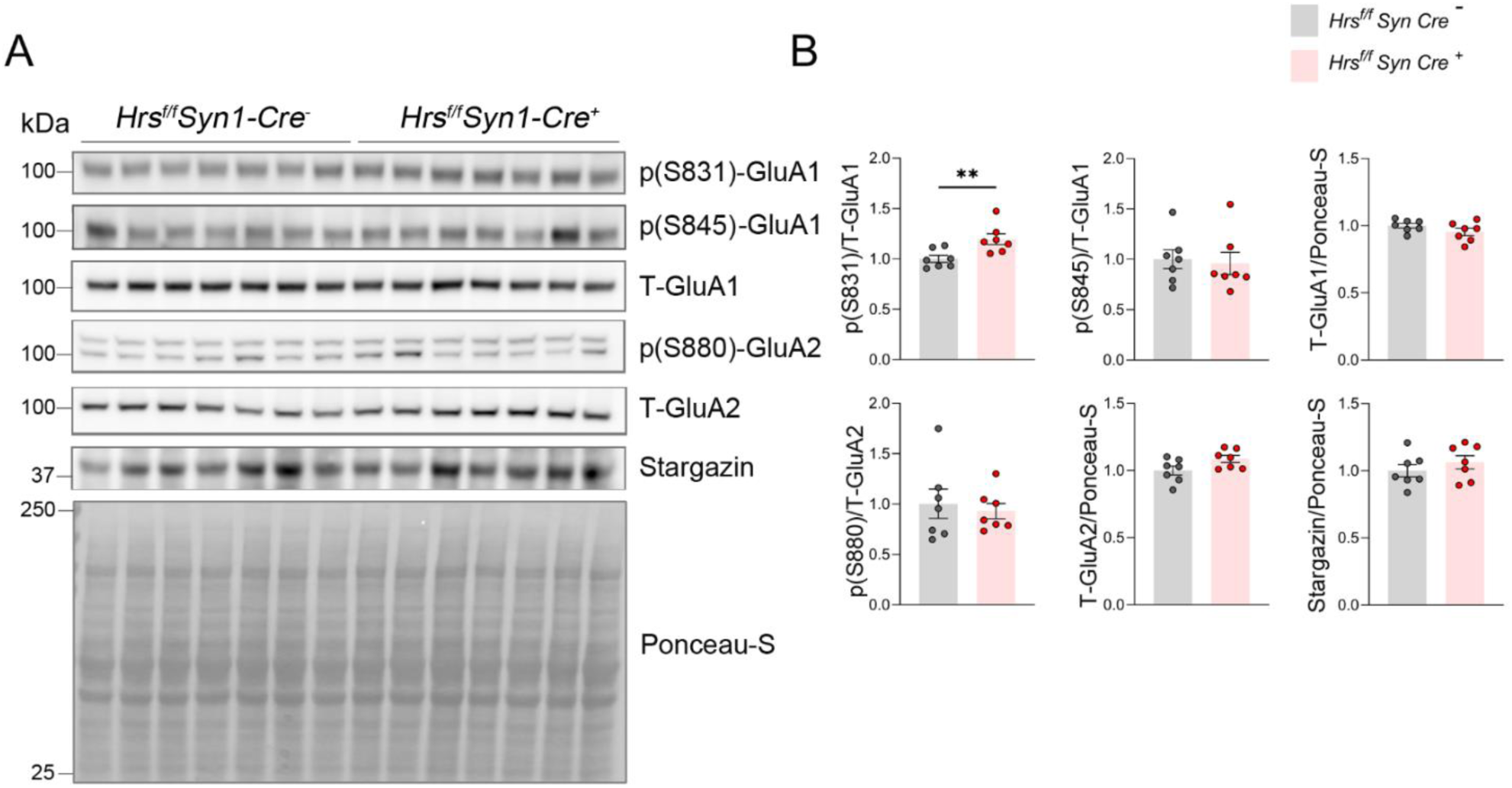
Neuronal Hrs depletion modestly enhances the phosphorylation of AMPAR subunit GluA1 at S831. (A) Western blots of cortical lysates from *Hrs^f/f^Syn1-Cre^-/-^* and *Hrs^f/f^Syn1-Cre^+/-^* mice for GluA1 and GluA2 subunits and their phosphorylation sites (n = 7 per group). (B) Densitometric quantification of western blot signals. Mean ± SEM. Unpaired two-tailed Student’s *t*-test; ** *p* < 0.01.

**Figure S9.**
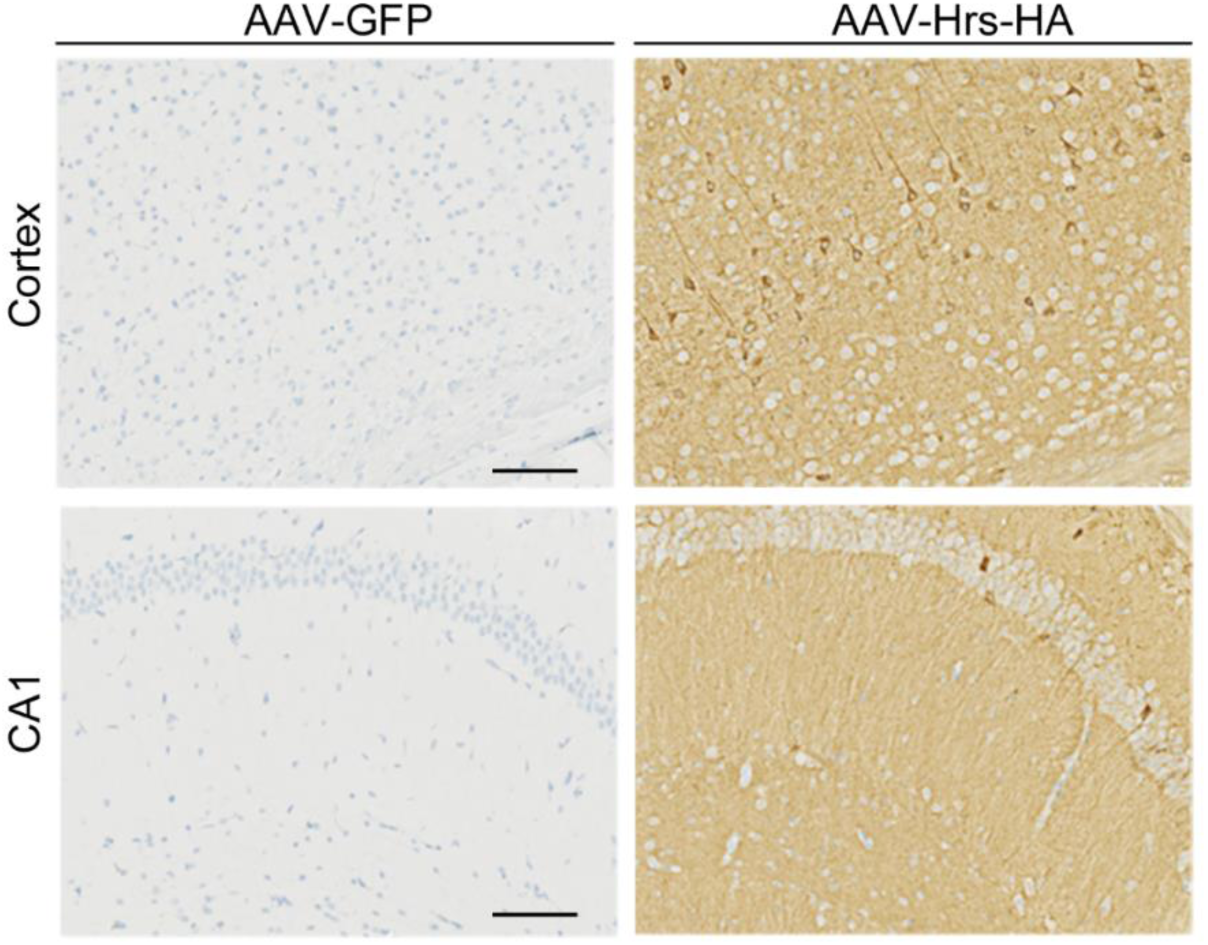
Robust neuronal expression of Hrs-HA in the cortex and hippocampus following AAV-Syn1-Hrs-HA transduction. Immunohistochemical detection of Hrs-HA in cortex and hippocampus (CA1) of adult WT mice three weeks after intravenous injection of AAV-Hrs-HA or control AAV-GFP (both of which express their transgene under the hSyn1 promoter). Strong HA immunoreactivity confirms robust neuronal Hrs overexpression. Scale bar = 100 µm.

**Figure S10.**
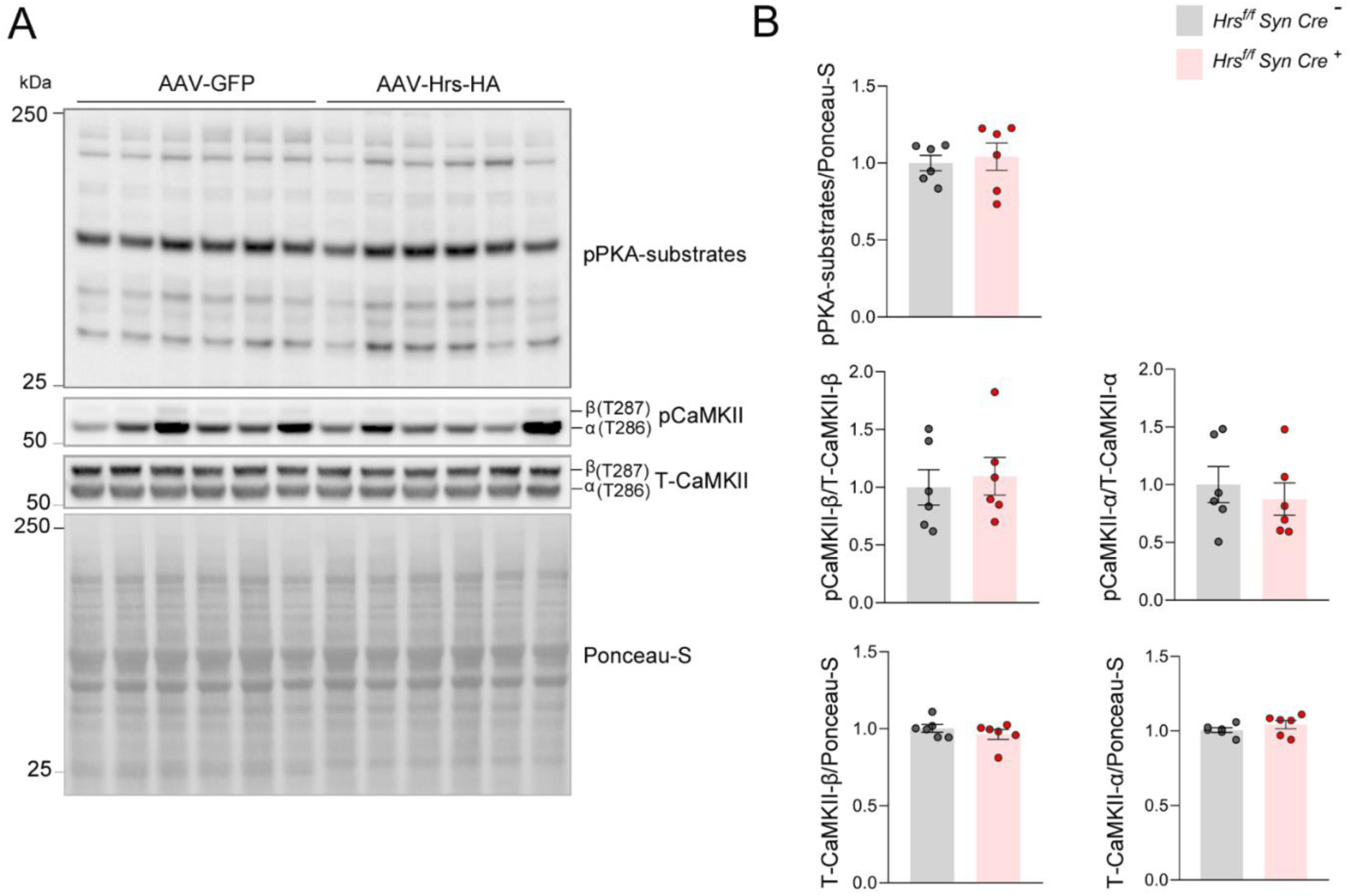
Neuronal Hrs overexpression in AAV-transduced mice did not affect the levels of phosphorylated or total CaMKII or phosphorylated PKA substrates. (A) Western blots of cortical lysates from adult WT mice injected with AAV-GFP or AAV-Hrs-HA (n = 6 per group), showing levels of phosphorylated PKA substrates (pPKA substrates) and phosphorylated and total CaMKII (p-T286) proteins. (B) Quantification of protein levels. Mean ± SEM. Unpaired two-tailed t-test.

